# Increased association between Epstein-Barr virus EBNA2 from type 2 strains and the transcriptional repressor BS69 restricts B cell growth

**DOI:** 10.1101/464131

**Authors:** Rajesh Ponnusamy, Ritika Khatri, Paulo B. Correia, Erika Mancini, Paul J. Farrell, Michelle J. West

## Abstract

Natural variation separates Epstein-Barr virus (EBV) into type 1 and type 2 strains. Type 2 EBV is less transforming *in vitro* due to sequence differences in the EBV transcription factor EBNA2. This correlates with reduced activation of the EBV oncogene LMP1 and some cell genes. Transcriptional activation by type 1 EBNA2 can be suppressed through the binding of two PXLXP motifs in its transactivation domain (TAD) to the dimeric coiled-coil MYND domain (CC-MYND) of the BS69 repressor protein (ZMYND11). We identified a third conserved PXLXP motif in type 2 EBNA2. We found that type 2 EBNA2 peptides containing this motif bound BS69_CC-MYND_ efficiently and that the type 2 EBNA2_TAD_ bound an additional BS69_CC-MYND_ molecule. Full-length type 2 EBNA2 also bound BS69 more efficiently in pull-down assays. Molecular weight analysis and low-resolution structures obtained using small-angle X-ray scattering showed that three BS69_CC-MYND_ dimers bound two molecules of type 2 EBNA2_TAD_, in line with the dimeric state of full-length EBNA2 *in vivo*. Importantly, mutation of the third BS69 binding motif in type 2 EBNA2 improved B-cell growth maintenance. Our data indicate that increased association with BS69 restricts growth promotion by EBNA2 and may contribute to reduced B-cell transformation by type 2 EBV.

**Author summary:** Epstein-Barr virus (EBV) drives the development of many human cancers worldwide including specific types of lymphoma and carcinoma. EBV infects B lymphocytes and immortalises them, thus contributing to lymphoma development. The virus promotes B lymphocyte growth and survival by altering the level at which hundreds of genes are expressed. The EBV protein EBNA2 is known to activate many growth-promoting genes. Natural variation in the sequence of EBNA2 defines the two main EBV strains: type 1 and type 2. Type 2 strains immortalise B lymphocytes less efficiency and activate some growth genes poorly, although the mechanism of this difference is unclear. We now show that sequence variation in type 2 EBNA2 creates a third site of interaction for the repressor protein (BS69, ZMYND11). We have characterised the complex formed between type 2 EBNA2 and BS69 and show that three dimers of BS69 form a bridged complex with two molecules of type 2 EBNA2. We demonstrate that mutation of the additional BS69 interaction site in type 2 EBNA2 improves its growth-promoting function. Our results therefore provide a molecular explanation for the different B lymphocyte growth promoting activities of type 1 and type 2 EBV. This aids our understanding of immortalisation by EBV.

## Introduction

Epstein-Barr virus (EBV) is a ubiquitous γ-herpesvirus that immortalises human B lymphocytes to establish a lifelong persistent infection that is usually harmless. Delayed primary EBV infection can however give rise to infectious mononucleosis. EBV is also associated with the development of malignancies that include Burkitt’s (BL), Hodgkin’s, diffuse large B cell and post-transplant lymphoma and nasopharyngeal or gastric carcinoma. EBV expresses nine latent proteins in *in vitro* infected lymphoblastoid cell lines (LCLs), including 6 Epstein-Barr nuclear antigens (EBNA1, 2, 3A, 3B, 3C and leader protein) and 3 latent membrane proteins (LMP1, 2A, 2B). The EBNA2 transcription factor is one of five of these latent genes essential for B cell transformation (1). EBNA2 functions as the master regulator of EBV latent gene transcription and activates numerous cell genes that control B cell growth and survival (2). It cannot however bind to DNA directly and hijacks cell DNA binding proteins e.g. RBPJ (RBPJκ, CBF1) and EBF1 to target viral and cell gene regulatory elements (2, 3). Although EBNA2 binding sites are close to gene promoters in the viral genome, in the B cell genome they are mostly found at enhancer elements and EBNA2 has been shown to promote enhancer-promoter interactions (4-6). EBNA2 activates transcription through interactions between its acidic transactivation domain (TAD) and histone acetyl transferases, ATP-dependent remodellers and components of the preinitiation complex (7-13).

EBV genome sequences worldwide separate into two main strains (type 1 and type 2) based on differences in the EBNA2 and EBNA3A, 3B and 3C genes (14-18). Type 2 strains are less efficient at immortalising resting B cells *in vitro* than type 1 strains (19). This phenotype is determined by sequence variation in EBNA2 since complementation of an EBNA2 defective virus with type 1 EBNA2 but not type 2 EBNA2 supports efficient primary B cell immortalisation (1). Consistent with its reduced primary B cell transforming function, type 2 EBNA2 cannot complement loss of type 1 EBNA2 function to maintain the growth of lymphoblastoid cell lines (20). Amino acids responsible for the differences in B cell growth maintenance between type 1 and type 2 EBNA2 were mapped to the C-terminal region of EBNA2 (20). Surprisingly, a single amino acid (aspartate 442 of type 1 EBNA2) in the TAD appears to be a key determinant of B cell growth maintenance by type 1 EBNA2. Replacing the serine that occurs at the equivalent position in type 2 EBNA2 (amino acid 409 of type 2 EBNA2) with aspartate (mutant S442D) confers efficient growth maintenance function (21). Type 2 EBNA2 has reduced ability to activate expression of the EBV oncogene LMP1 and a small number of cellular genes e.g. CXCR7 (22). These differences in gene activation could underlie the reduced B cell growth maintenance and transforming function of type 2 EBNA2, although the mechanism involved and the role played by the single aspartate residue is unclear. Part of the mechanism may involve (or result in) reduced binding of type 2 EBNA2 to the LMP1 promoter and cell gene regulatory elements (21). EBNA2 binding sites at genes activated less efficiently by type 2 EBNA2 are enriched for composite binding motifs for ETS and IRF transcription factors (ETS and IRF composite element; EICE), implicating ETS/IRF family members in the gene specificity of the observed effects (21).

Despite a clear deficiency in the immortalising and B cell growth maintenance properties of type 2 EBNA2 *in vitro*, no specific differences in disease association have been reported to date for type 1 and type 2 EBV. Interestingly, although the outgrowth of immortalised cells is less efficient and much slower in primary B cell cultures infected with type 2 EBV (19, 20), the LCLs that are eventually established from type 2 viruses proliferate at similar rates to type 1 LCLs. Type 1 and 2 LCLs also show equivalent expression of LMP1 and CXCR7 (20). Over extended periods of time, it is therefore possible to select for immortalised cells infected with type 2 EBV that have the required levels of expression of these genes to support their long term proliferation. *In vivo* other factors may create an environment that helps support B cell immortalisation by type 2 EBV.

New research also suggests that type 2 EBV may use alternative approaches to persist *in vivo*. Type 2 EBV has the unique capacity to infect T cells in culture and is detected in T cells from healthy infants from Kenya, indicating that T cell infection may form part of a natural type 2 EBV infection (23, 24). Recent work also showed that a type 2 EBV strain was able to infect both B cells and T cells in humanised mice (25). Mice infected with type 2 EBV developed tumours that resembled the diffuse large B-cell lymphomas that also developed in mice infected with type 1 EBV, confirming the tumorigenic potential of a persistent type 2 EBV infection, once established (25).

BS69 (ZMYND11) is a multi-domain chromatin-associated repressor protein that suppresses transcription elongation, regulates pre-mRNA processing and has tumour suppressor function (26, 27). The BS69 gene undergoes chromosomal translocation in minimally differentiated myeloid leukaemia leading to the expression of a BS69-MBTD1 fusion protein (28). BS69 contains three histone reader domains in its N-terminal region; a plant homeodomain, a bromodomain and a PWWP domain. The tandemly-arranged bromodomain and PWWP domain bind to histone H3 or the variant histone H3.3 when trimethylated on lysine K36 (27, 29). BS69 also contains a coiled-coil (CC) dimerisation domain adjacent to a MYND domain in its C terminus. BS69 binds to a number of chromatin modifying enzymes (BRG1, HDAC1, EZH2) and transcription factors (adenovirus E1a, c-Myb, ETS2, E2F6, and the Myc-associated MGA protein) and inhibits transcription factor activation function (30-34). The BS69 MYND domain binds to E1a and MGA through a PXLXP motif (34).

BS69 has also been shown to interact with the TAD of type 1 EBNA2 through two PXLXP motifs and to restrict EBNA2 transcriptional activation function (34, 35). The structure of the dimeric CC-MYND domain of BS69 bound to two peptides encompassing sequences from one of the EBNA2 PXLXP motifs (motif 1) has been solved (35). Based on this three-dimensional structure, a BS69 dimer was predicted to interact with the two adjacent PXLXP motifs in type 1 EBNA2. Interestingly the amino acid implicated in the type-specific differences in growth maintenance observed for type 1 and type 2 EBNA2 (amino acid 442 in type 1 EBNA2) (21) lies immediately adjacent to the second BS69 binding motif.

We set out to determine whether sequence differences between type 1 and type 2 EBNA2 affect BS69 binding. We hypothesised that type 2 EBNA2 would show increased binding to BS69 and that this would impair its gene activation and growth maintenance function. We initially examined the impact of type-specific differences in EBNA2 amino acid 442 on BS69 binding. We also identified a third PXLXP BS69 binding motif in type 2 EBNA2, so we examined whether the presence of this extra motif resulted in the interaction of additional molecules of BS69 with type 2 EBNA2. We found that amino acid 442 did not affect BS69 binding, but influenced the conformation of the TAD, potentially affecting binding of other transcriptional regulators. We also demonstrated that the third PXLXP motif in type 2 EBNA2 was responsible for the binding of an additional BS69 dimer. Importantly, mutation of the third PXLXP BS69 binding motif in full length type 2 EBNA2 restored B cell growth maintenance function indicating that increased BS69 binding is responsible for impaired type 2 EBNA2 function.

## Results

### Sequence differences near type 2 EBNA2 BS69 binding motif 2 do not increase BS69 binding

Previous studies demonstrated that differences in a single amino acid in the TAD between type 1 and type 2 EBNA2 determined the ability of EBNA2 to maintain the growth of an EBV-infected LCL (21). This amino acid (located at position 442 in the EBNA2 sequence from the prototypical type 1 B95-8 strain of EBV) is conserved as aspartate in type 1 strains and as serine (at the corresponding position of 409) in type 2 strains. In type 1 EBNA2, aspartate 442 is located immediately adjacent to a previously identified binding motif for the cell transcriptional repressor BS69 (motif 2) that fits the PXLXP consensus (PILFP_437-441_)(34). In the TAD of type 2 EBNA2, the PXLXP motif is conserved (PFLFP_404-408_) and is flanked by serine 409 (Figure 1A).

**Figure 1.**
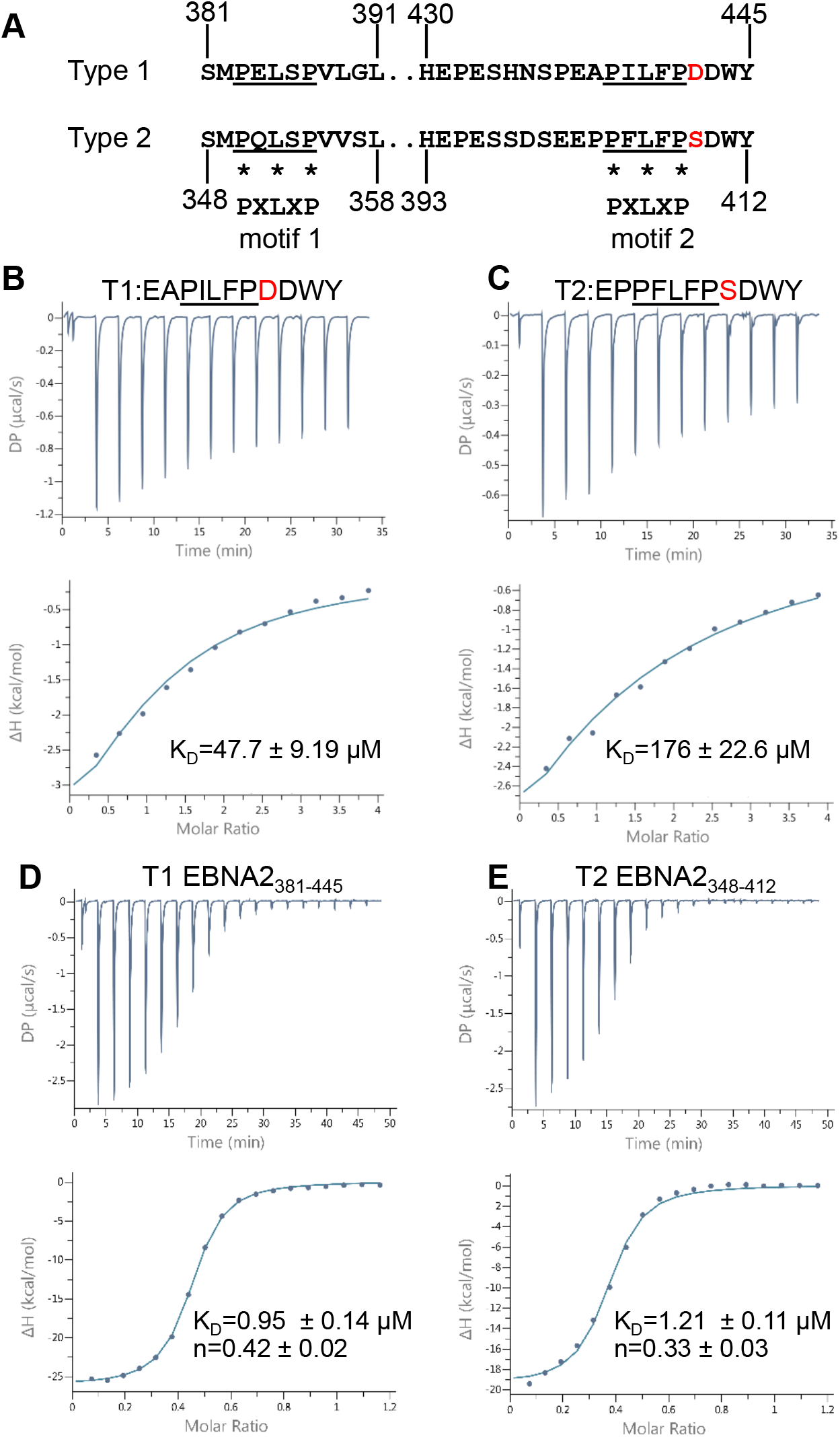
Isothermal titration calorimetry analysis of the interaction of type 1 and type 2 EBNA2 peptides and polypeptides containing BS69 binding motif 2 with BS69_CC-MYND_. (A) Amino acid sequence of the regions of type 1 (B95-8) and type 2 (AG876) EBNA2 containing BS69 binding motifs 1 and 2 (underlined). Asterisks show the positions of the P, L and P amino acids in the PXLXP binding motif. Aspartate 442 in type 1 EBNA2 and the corresponding serine 409 in type 2 EBNA2 adjacent to motif 2 are shown in red. (B) Isothermal titration calorimetry (ITC) analysis of type 1 EBNA2 motif 2 peptide binding to BS69_CC-MYND_. The upper panel shows heat peak data as Differential Power (DP) versus time and the lower panel shows ΔH (derived from integration of the heat peak intensities) plotted against the BS69_CC-MYND_/EBNA2 molar ratio (based on monomer concentrations). Titrations were performed using a series of 13 2.0 μl injections of 1 mM EBNA2 peptide and 0.1 mM BS69_CC-_ MYND in the cell. The solid line shows the best fit using a one-site (one event) binding model with the n value fixed to 1. The dissociation constant (K_D_) displayed shows the mean ± standard deviation obtained from 3 independent experiments (C) ITC analysis of the binding of the type 1 EBNA2 polypeptide T1 EBNA2_381-445_ (0.6 mM) to BS69_CC-MYND_ using 19 2.0 μl injections of 0.1 mM EBNA2 polypeptide. The n value indicates the stoichiometry of BS69_CC-_ MYND/EBNA2 binding calculated for monomeric proteins and shows the mean ± standard deviation from 3 independent experiments. (D) ITC analysis of the binding of the type 2 EBNA2 polypeptide T2 EBNA2_348-412_ (0.3 mM) to BS69_CC-MYND_ (0.1 mM) as in C.

We hypothesised that the impaired gene activation and growth maintenance properties of type 2 EBNA2 may be the result of increased binding to BS69 as a result of the aspartate to serine amino acid difference adjacent to BS69 binding motif 2. We therefore tested whether a BS69 binding motif 2 peptide from type 2 EBNA2 showed enhanced binding to BS69 compared to a motif 2 peptide from type 1 EBNA2. We used isothermal titration calorimetry (ITC) to determine the affinity of peptide binding to the C-terminal region of BS69 (amino acid 480-602) comprising the CC-MYND domain that we expressed and purified from *E.coli* (Figure 1B and 1C). In contrast to our hypothesis, we found that the type 2 EBNA2 motif 2 peptide bound to BS69_CC-MYND_ with reduced affinity (K_D_=176 μM) compared to the corresponding peptide from type 1 EBNA 2 (K_D_=47.7 μM) (Figure 1B and 1C and Supplementary Table S1). The affinity of binding of the type 1 EBNA 2 motif 2 peptide to BS69_CC-MYND_ was very similar to the previously reported K_D_ of 35 μM (35). The difference in binding between type 1 and type 2 EBNA2 motif 2 peptides could be influenced by both the aspartate to serine change and differences in two other amino acids present in the sequence (Figure 1B and 1C).

An additional BS69 PXLXP binding motif previously identified in type 1 EBNA2 (motif 1) located N-terminal to motif 2 is also present in type 2 EBNA 2 (Figure 1A). The BS69_CC-MYND_ dimer binds a type 1 EBNA2 polypeptide containing both motif 1 and motif 2 with high affinity and the structure of BS69 dimer could accommodate binding to both motifs simultaneously (35). We therefore tested whether sequence differences in type 2 EBNA2 (including the aspartate to serine change) affected the binding of a region of EBNA2 containing both motif 1 and motif 2 to BS69. Type 1 EBNA2_381-445_ and type 2 EBNA2_348-412_ were expressed and purified from *E.coli* and their interaction with BS69_CC-MYND_ examined using ITC. Consistent with previous reports (35) we found that type 1 EBNA2_381-445_ bound to BS69_CC-MYND_ with high affinity (K_D_=0.95 μM) likely due to the high avidity of interaction with two binding sites (Figure 1D). In the context of this larger region of EBNA2 we found very little difference in the affinity of type 2 EBNA2 binding to BS69_CC-MYND_ (K_D_=1.21 μM) (Figure 1E). In addition to measuring binding affinities, ITC data can also be used to calculate binding stoichiometry (n) which can be visualised as the molar ratio at the mid (inflection) point of the sigmoidal binding curve. We titrated EBNA2 polypeptides into a cell containing BS69_CC-MYND,_ so the stoichiometry values we obtained indicate the molar ratio at which the EBNA2 polypeptide saturates the available sites in BS69_CC-MYND_ monomers. Consistent with the presence of two BS69 binding sites in the EBNA2 polypeptides, we obtained n values of 0.42 and 0.33 for type 1 and type 2 EBNA2: BS69_CC-MYND_ binding, respectively (Figure 1D and 1E). These approximate to the expected molar ratio of 0.5 taking into consideration some margin of error in n value determination by ITC, which is heavily influenced by the accuracy of protein concentrations and the proportion of ‘active’ protein in the sample.

We conclude that the aspartate 442 to serine amino acid difference between type 1 and type 2 EBNA2 does not affect the binding of BS69 to the TAD of type 2 EBNA2 in these assays. Our data also indicate that additional sequence differences in and around BS69 binding motifs 1 and 2 in type 2 EBNA2 do not influence the binding of the BS69_CC-MYND_ dimer to this region of the protein.

### Type 2 EBNA2 contains a third BS69 binding site

During the course of our study we also identified a third potential BS69 binding site in the type 2 EBNA2 TAD (Figure 2A). In type 2 EBNA2, a sequence that is an exact match to the PXLXP BS69 consensus binding motif is present C-terminal to motif 2 (PTLEP_414-418_). In type 1 EBNA2 the corresponding region has an isoleucine in place of the leucine residue (PSIDP_447-451_). To determine whether these regions of EBNA2 also interact with BS69, we performed ITC experiments using type 1 and type 2 EBNA2 peptides (Figure 2B and 2C). We were not able to detect any binding of the type 1 EBNA2 peptide encompassing this region (T1 EBNA2_445-455_) to BS69_CC-MYND_, underscoring the importance of the central leucine in the PXLXP motif for the BS69 interaction (Figure 2B). In contrast, a peptide from the corresponding region of type 2 EBNA2 (T2 EBNA2_412-422_) interacted with BS69 with a K_D_=219 μM (Figure 2C). The affinity of interaction with this new motif (that we named motif 3) is weaker than the interaction we observed for type 1 or type 2 EBNA2 motif 2 (Figure 1). To determine the impact of motif 3 on the interaction of type 2 EBNA2 with BS69_CC-MYND_ in the presence of the two other BS69 binding motifs, we expressed and purified a larger type 2 EBNA2 polypeptide containing motif 1, 2 and 3 (T2 EBNA2_348-422_) for use in ITC. For comparison, we also analysed the binding of the corresponding larger region of type 1 EBNA2 (T1 EBNA2_381-455_). We found that inclusion of the additional C-terminal amino acids had little impact on the affinity of binding of type 1 EBNA2 to BS69_CC-MYND_ or the stoichiometry of binding (compare Figure 2D and Figure 1D) (Supplementary Table S2). In contrast, for type 2 EBNA2, we observed a change in the stoichiometry of binding from 0.33 when motif 1 and 2 were present (T2 EBNA2_348-412)_ to 0.15 when motif 1, 2 and 3 were present (T2 EBNA2_348-422_) (compare Figure 2E and 1E). This is consistent with the presence of an additional BS69 binding site and indicates the interaction of T2 EBNA2_348-422_ with an additional BS69_CC-MYND_ molecule. Perhaps surprisingly, we did not observe an increase in the affinity of binding of the longer type 2 EBNA2 polypeptide to BS69_CC-MYND_ (compare Figure 2E and 1E). Nonetheless the recruitment of more BS69 to type 2 EBNA2 could be physiologically relevant for the function of type 2 EBNA2 as a transcriptional activator. To confirm that the observed change in binding stoichiometry was due to the presence of motif 3 in the type 2 EBNA2 polypeptide, we analysed the binding of T2 EBNA2_348-422_ with motif 3 mutated from PTLEP to ATAEA (T2 EBNA2_348-422_ motif 3 mt). We found that mutation of motif 3 altered the stoichiometry of binding to BS69_CC-MYND_ from 0.15 to 0.30, consistent with the loss of a BS69 binding motif (Figure 2F). This is similar to the value obtained for the type 2 EBNA2 polypeptide containing only motif 1 and motif 2 (T2 EBNA2_348-412_)(Figure 1E). We conclude that type 2 EBNA2 contains an additional binding site for BS69 that is not present in type 1 EBNA2.

**Figure 2.**
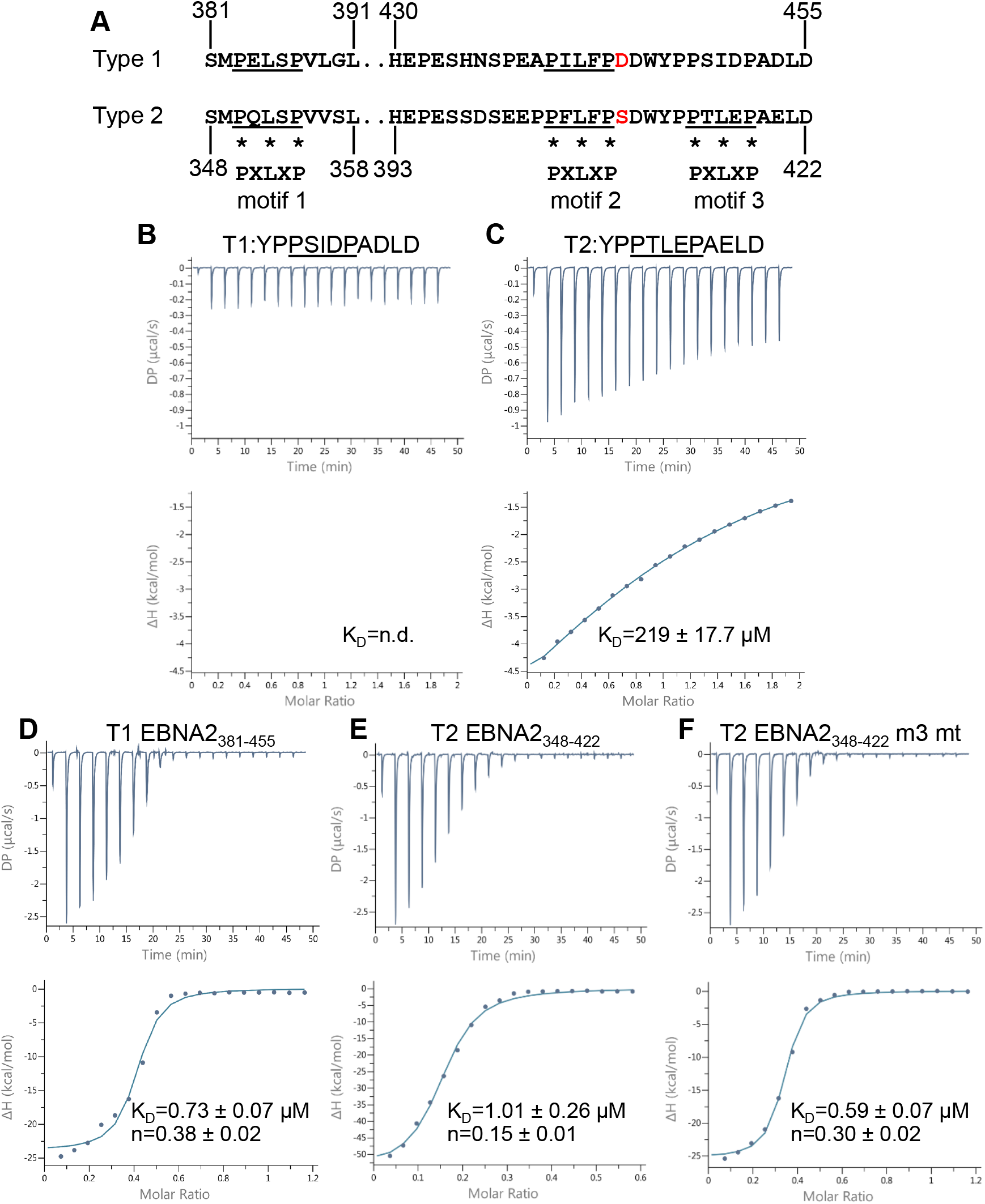
Isothermal titration calorimetry analysis of the interaction of type 1 and type 2 EBNA2 peptides and polypeptides containing BS69 binding motif 3 with BS69_CC-MYND_. (A) Amino acid sequence of the regions of type 1 (B95-8) and type 2 (AG876) EBNA2 containing BS69 binding motifs 1, 2 and 3 (underlined). Asterisks show the positions of the P, L and P amino acids in the PXLXP binding motif. (B) Isothermal titration calorimetry (ITC) analysis of type 1 EBNA2 motif 3 peptide binding to BS69_CC-MYND_ (no binding was detected so the K_D_ could not be determined (n.d.). (C) Isothermal titration calorimetry (ITC) analysis of type 2 EBNA2 motif 3 peptide binding to BS69_CC-MYND_ using 19 injections of 2.0 μl EBNA2 peptide. Data are displayed and analysed as in Figure 1. (C) ITC analysis of the binding of type 2 EBNA2 motif 3 peptide binding to BS69_CC-MYND_ as in B. (D) ITC analysis of the binding of the type 1 EBNA2 polypeptide EBNA2_381-455_ to BS69_CC-MYND._ (E) ITC analysis of the binding of the type 2 EBNA2 polypeptide T2 EBNA2_348-422_ to BS69_CC-MYND_. (F) ITC analysis of the binding of the type 2 EBNA2 polypeptide T2 EBNA2_348-422_ with BS69 binding motif 3 mutated from PTLEP to ATAEA.

### Three BS69 CC-MYND dimers bridge two molecules of type 2 EBNA2

To further examine whether type 2 EBNA2 could form higher-order complexes with the BS69 CC-MYND domain that are larger than type 1 EBNA2, we examined the properties of BS69-EBNA2 complexes using size exclusion chromatography (SEC). Consistent with complex formation, when pre-incubated with BS69_CC-MYND,_ both T1 EBNA2_381-455_ and T2 EBNA2_348-422_ polypeptides migrated through the size exclusion column faster and eluted at a lower elution volume compared to the migration of each component individually (Figure 3A). In line with the binding of additional BS69_CC-MYND_ molecules to T2 EBNA2_348-422_ and the formation of higher molecular weight complexes, we found that type 2 EBNA2 complexes eluted at a lower volume than type 1 EBNA2 complexes (Figure 3A). SDS-PAGE of SEC column fractions confirmed the presence of BS69_CC-MYND_ and EBNA2 in the higher molecular weight complexes (Figure 3B). Note that both type 1 and type 2 EBNA2 polypeptides migrate anomalously on SDS-PAGE gels and not at their predicted molecular weights (MW) of 7.9 and 8.1 kDa respectively, likely due to their high proline content (Figure 3B). They are however pure and resolve as single species on gel filtration columns (Figure 3A).

**Figure 3.**
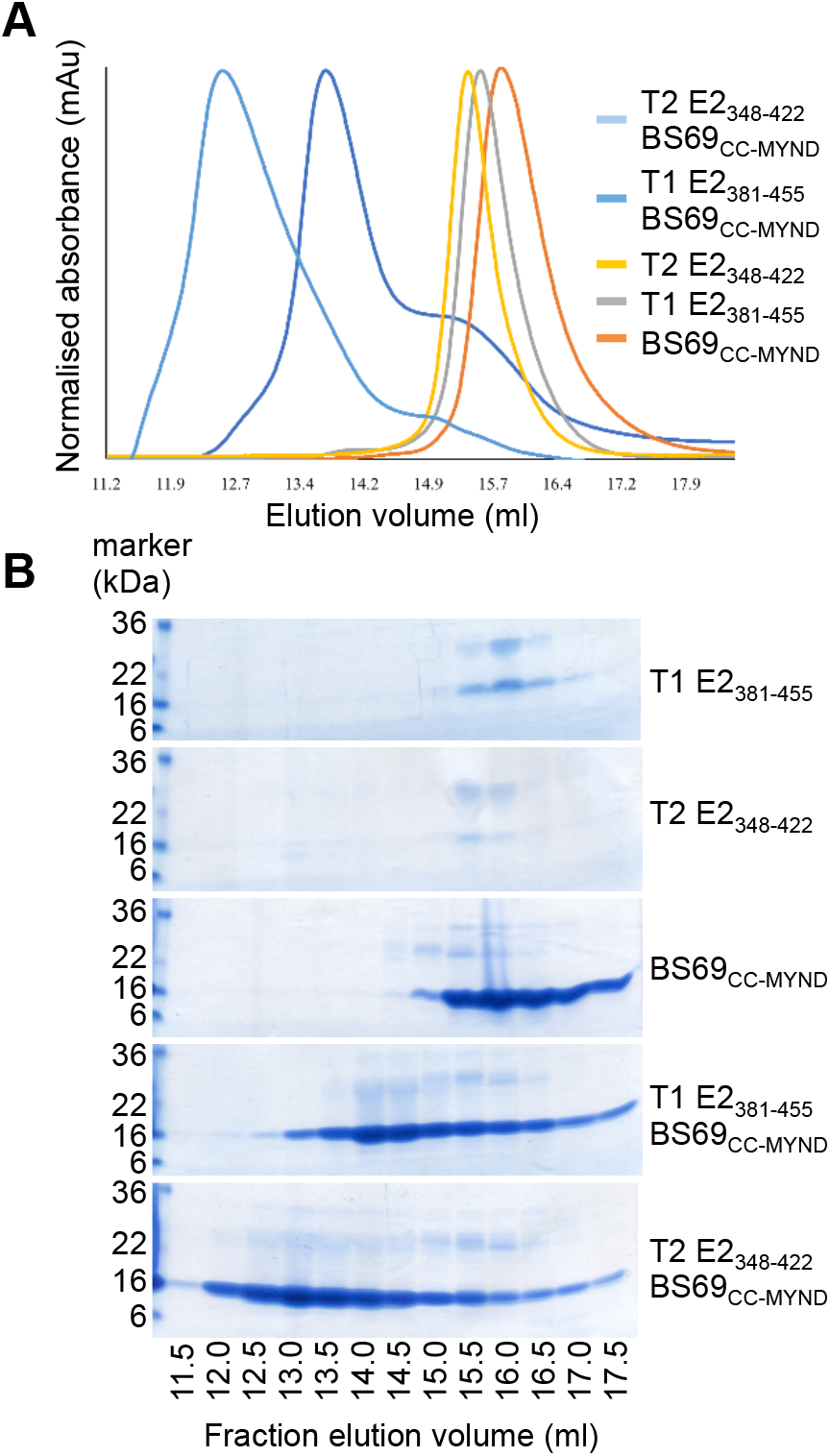
Solution state analysis of EBNA2 and BS69_CC-MYND_ complexes. (A) Size exclusion chromatography of individual type 1 and type 2 EBNA2 polypeptides, BS69_CC-MYND_ and EBNA2-BS69_CC-MYND_ complexes. Absorbance was normalized to the type2 EBNA2-BS69 complex (highest absorbance) using UNICORN software. (B) Samples from the indicated fractions were analysed by SDS-PAGE followed by Coomassie staining.

Because migration in SEC is influenced by both size and shape and BS69_CC-MYND_ has an elongated structure due to the CC domain, we were unable to determine the MW of BS69-EBNA2 complexes accurately using SEC. In order to obtain more accurate MW information that would allow us to determine the number of molecules of BS69_CC-MYND_ and EBNA2 present in type 1 and type 2 complexes, we used SEC with multi-angle light scattering (SEC-MALS) (Table 1). SEC-MALS gave MWs for T1 EBNA2_381-455_ and T2 EBNA2_348-422_ that matched the theoretical MW of their monomeric forms and gave a MW for BS69_CC-MYND_ consistent with its dimeric state (Table 1). For the T1 EBNA2_381-455_-BS69_CC-MYND_ complex, SEC-MALS gave a MW of 62.3 kDa. Given that there are two binding sites for BS69 in the T1 EBNA2_381-455_ polypeptide, this figure most closely matches the MW of a complex containing two type 1 EBNA2 polypeptides and two BS69_CC-MYND_ dimers (theoretical MW of 76.7 kDa) rather than a single type 1 EBNA2 polypeptide with a one dimer of EBNA2 BS69_CC-MYND_ (theoretical MW of 38.3 kDa) (Table 1). For the BS69_CC-MYND_-T2 EBNA2_348-422_ complex, SEC-MALs gave a MW of 135 kDa consistent with the larger complex size observed in SEC (Table 1 and Figure 3A). Given the presence of three BS69 binding motifs in type 2 EBNA2, this MW most closely matches that of a complex containing three BS69_CC-MYND_ dimers and two type 1 EBNA2 polypeptides (theoretical MW of 107.5 kDa) (Table 1).

**Table 1.** Size exclusion chromatography and multiangle might scattering (MALS) determination of the molecular weight of individual type 1 and type 2 EBNA2 polypeptides, BS69_CC-MYND_ and EBNA2-BS69 complexes. Theoretical and experimentally determined molecular weights are shown. For EBNA2-BS69 complexes, the theoretical molecular weights of complexes containing different numbers of EBNA2 and BS69 molecules are shown in parentheses. The most likely solution state based the experimentally determined molecular weight is indicated for each sample.

|  | T1 E2<sub>381-455</sub> | T2 E2<sub>348-422</sub> | BS69<sub>CC-MYND</sub> | T1 E2<sub>381-455</sub><br>BS69<sub>CC-MYND</sub> | T2 E2<sub>348-422</sub><br>BS69<sub>CC-MYND</sub> |
| --- | --- | --- | --- | --- | --- |
| Theoretical MW (kDa) | 7.9 (monomer) | 8.1 (monomer) | 15.2 (monomer) | 38.3 (1:2)<br>76.7 (2:4)<br>153.2 (4:8) | 53.7 (1:3)<br>107.5 (2:6)<br>215 (4:12) |
| SEC-MALS MW (kDa) | 7.5 | 8.2 | 35.4 | 62.3 | 135 |
| Solution state | T1 E2 monomer | T2 E2 monomer | BS69<sub>CC-MYND</sub> dimer | 2 x T1 E2<br>2 x BS69<sub>CC-MYND</sub> dimers | 2 x T2 E2<br>3 x BS69<sub>CC-MYND</sub> dimers |

Because of the discrepancies in the theoretical and experimentally determined MWs for BS69_CC-MYND_-EBNA2 complexes, we also used small-angle-X-ray scattering (SAXS) to obtain information on the shape and size of these complexes in solution. Initially we used SEC-SAXS to analyse each polypeptide individually. We used a Kratky representation to visualize features of the scattering profiles obtained for T1 EBNA2_381-45,_ T2 EBNA2_348-422_ and BS69_CC-MYND_ individually to identify the folding state of the polypeptides in solution. The absence of a bell-shaped curve with a well-defined maximum for both EBNA2 polypeptides indicates that they are natively unfolded in solution (Supplementary Figure S1). The bell-shaped curve obtained for the BS69_CC-MYND_ dimer indicates that it is folded in solution as expected from the crystal structure (35). Three-dimensional models were created for the individual polypeptides by *ab initio* shape determination. For BS69_CC-MYND_ a solution structure consistent with the coiled-coil dimer structure determined by X-ray crystallography was obtained (35)(Supplementary Figure S2). For the EBNA2 polypeptides, solution structures consistent with flexible unfolded peptide chains were obtained (Supplementary Figure S2). SAXS analysis of BS69_CC-MYND_ pre-mixed with either type 1 or type 2 EBNA2 polypeptides gave a larger Porod volume (directly related to MW) compared to the individual proteins, consistent with complex formation (Supplementary Table S3). An *ab initio* dummy atom model was generated for the type 1 EBNA2-BS69_cc-MYND_ complex and this fitted well to the experimental SAXS data (χ^2^ of 1.4) (Figure 4A). The three-dimensional model generated by *ab initio* shape determination for the type 1 EBNA2-BS69_cc-MYND_ complex indicated that the complex has a large elongated shape with a volume of 166 nm3 and a maximum dimension (Dmax) of 138 Å (Figure 4B). This three-dimensional model could accomodate two BS69_CC-MYND_ dimer structures that were manually docked into the SAXS envelope. The additional space at the bottom of the model was allocated to the model solution structures of two type 1 EBNA2 polypeptides (Figure 4B). This docked structural model for the type 1 EBNA2-BS69_CC-MYND_ complex was then fitted to the experimental scattering data and gave a reasonable χ^2^ value of 2.56. For comparison a structural model where only one BS69_CC-MYND_ dimer and a single type 1 EBNA2 polypeptide were docked into the SAXS envelope was created but this alternative model gave a worse fit to the experimental data (Supplementary Figure S3). An *ab initio* dummy atom model was then generated for the type 2 EBNA2-BS69_CC-MYND_ complex and this fitted well to the experimental SAXS data (χ^2^ of 1.0) (Figure 4C). The three-dimensional model created by *ab initio* shape determination for the type 2 EBNA2 complex had a larger volume (239 nm^3^) and maximum dimension (145 Å) than the type 1 EBNA2 complex model (Figure 4D). The type 2 EBNA2 model could accommodate the docking of three BS69_CC-MYND_ dimer structures along with two type 2 EBNA2 polypeptides and this structural model gave a good fit to the experimental data (χ^2^ of 1.44) (Figure 4C and D). In comparison a docked model containing two BS69_CC-MYND_ dimers and a single type 2 EBNA2 polypeptide gave a worse fit to the experimental data (Supplementary Figure S3).

**Figure 4.**
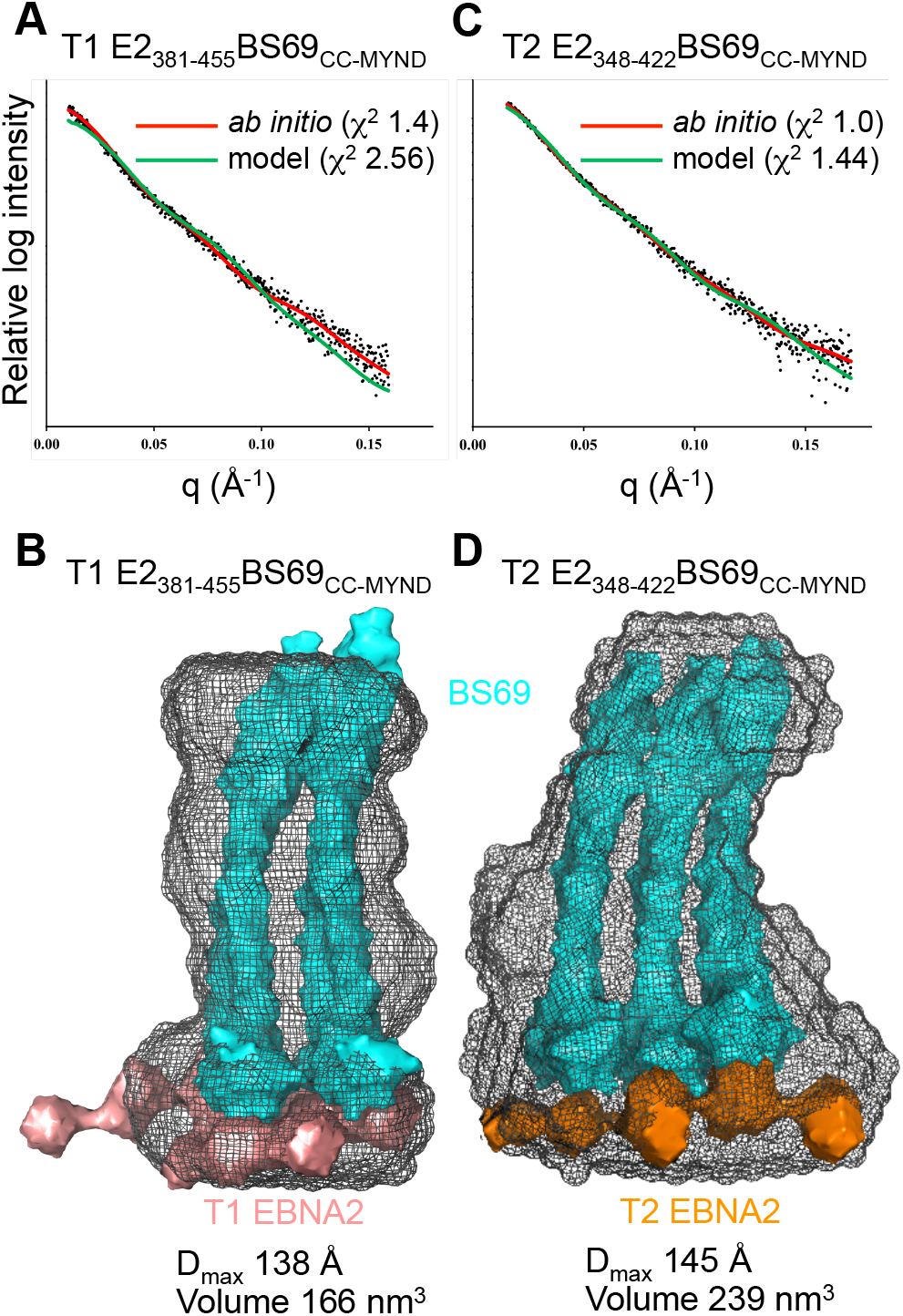
Solution structure of EBNA2-BS69_CC-MYND_ complexes determined by SAXS. (A) SAXS scattering data for type 1 EBNA2_381-455_-BS69_CC-MYND_ (black dots) fitted to the *ab initio* DAMMIN dummy atom model (red line). SAXS scattering data fitted to the docked structural complex shown in B (green) with the χ^2^ determined by FoXS. Plots show relative log intensity vs scattering vector (q). (B) Solution structure of type 1 EBNA2_381-455_-BS69_CC-MYND._ The SAXS envelopes (grey mesh) were generated by averaging 23 *ab-initio* models using the DAMMIF programme. The crystal structures of two BS69_CC-MYND_ dimers (PDB ID: 5HDA) are shown in cyan and were manually docked into the SAXS envelope along with the *ab-initio* dummy atom SAXS solution structures of two type 1 EBNA2_381-455_ polypeptides (salmon). The maximum dimension (D_max_) and volume were calculated using ScÅtter. (C) SAXS scattering data for type 2 EBNA2_348-422_-BS69_CC-MYND_ (black dots) fitted to the *ab initio* DAMMIN dummy atom model (red line). SAXS scattering data fitted to the docked structural complex shown in D (green) with the χ^2^ determined by FoXS. (D) Solution structure of type 2 EBNA2_348-422_-BS69_CC-MYND_ obtained as described in (B). The crystal structures of three BS69_CC-MYND_ dimers (PDB ID: 5HDA) are shown in cyan and were manually docked into the SAXS envelope along with the *ab-initio* dummy atom SAXS solution structures of two type 2 EBNA2_348-422_ polypeptides (orange).

Taken together our data indicate that BS69 forms higher order complexes with EBNA2 that involve the interaction of each MYND domain of the BS69 dimer with binding sites in two separate EBNA2 molecules. Rather than an *in vitro* artefact, this intermolecular ‘bridging’ interaction is consistent with the fact that EBNA2 forms dimers *in vivo*. Although the N-terminal regions of EBNA2 that mediate dimerisation (36) are absent in the EBNA2 polypeptides we examined in our interaction studies, our data indicate that BS69 may have the capacity to stabilise or enhance dimerisation between two EBNA2 molecules held together through their N-termini. Importantly, using multiple independent techniques, we also demonstrate that type 2 EBNA 2 interacts with an additional BS69_CC-MYND_ dimer.

### The serine to aspartate change in type 2 EBNA2 alters its binding characteristics

During the course of our ITC experiments we observed that binding data obtained using the longer EBNA2 polypeptides (T1 EBNA2_381-455_ and T2 EBNA2_348-422_) showed some deviation from curves fitted using the single binding event (‘one set of sites’) model (where binding to multiple sites cannot be detected as separate heat change events) (Figure 2D and 2E). This suggested that the mode of binding of these polypeptides to BS69_CC-MYND_ could involve more than one distinguishable binding event. To determine whether this was the case, we performed ITC experiments using an increased number of smaller injections of the EBNA2 polypeptide to obtain more data points for curve fitting (Figure 5). For T1 EBNA2_381-455_ the binding data did not fit well to curves generated using an alternative two binding event (‘two sets of sites’) model (χ^2^/degrees of freedom=0.56) (Figure 5A and Supplementary Table S2). This indicates that the deviation of T1 EBNA2_381-455_-BS69_CC-MYND_ binding data from fitted curves at low molar ratios was unlikely to be the result of a separate binding event (Figure 2D and 5A). In contrast, for T2 EBNA2_348-422_ the binding profiles obtained fitted well to curves generated using the two binding event model (χ^2^/degrees of freedom=0.23) (Figure 5B and Supplementary Table S2). This enabled the affinity of the two separate binding events to be determined, which were both in the nanomolar range (K_D1_=0.009 μM and K_D2_=0.091 μM). These data indicate that this region of type 2 EBNA2 may adopt a different conformation to type 1 EBNA2 when binding to BS69_CC-MYND._

**Figure 5.**
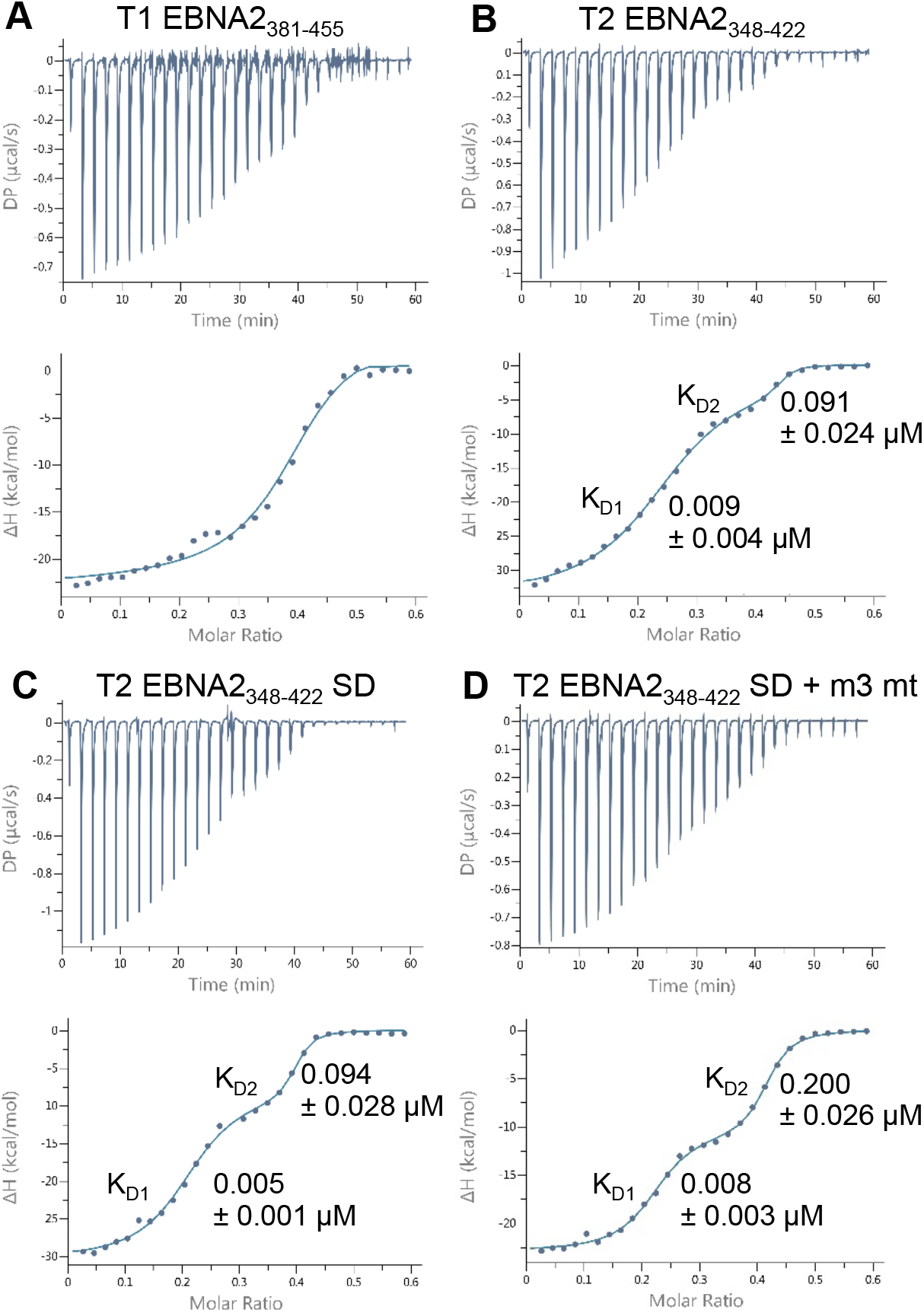
Isotheral titration calorimetry analysis of the interaction of type 1 and type 2 EBNA2 polypeptides with BS69_CC-MYND_ using an increased number of injections. (A) ITC analysis of the binding of the type 1 EBNA2 polypeptide EBNA2_381-455_ to BS69_CC-MYND._ Titrations were performed using a series of 29 injections of 1.3 μl 0.3 mM EBNA2 polypeptide and 0.1 mM BS69_CC-MYND_ in the cell. The solid line shows the best fit using a two-site (two event) binding model. The mean dissociation constant (K_D_) ± standard deviation from 3 independent experiments is shown for each binding event. (B) ITC analysis of the binding of the type 2 EBNA2 polypeptide T2 EBNA2_348-422_ to BS69_CC-MYND_ using 29 injections and fitting using a two-site (two event) binding model as in A. (C) ITC analysis of the binding of the type 2 EBNA2 polypeptide T2 EBNA2_348-422_ SD containing the serine 412 to aspartate mutation using 29 injections and fitting using a two-site (two event) binding model. (D) ITC analysis of the binding of the type 2 EBNA2 polypeptide T2 EBNA2_348-422_ with BS69 binding motif 3 mutated from PTLEP to ATAEA using 29 injections and fitting using a two-site (two event) binding model.

In further experiments we addressed the impact of changing the serine at position 409 in type 2 EBNA2 to the aspartate present at the equivalent position (aspartate 442) in type 1 EBNA2on BS69 binding in the context of the longer type 2 EBNA2 polypeptide containing three BS69 binding sites. To do this we expressed and purified a type 2 EBNA2 polypeptide with an S409D mutation (T2 EBNA2_348-422_ SD mutant). Interestingly, we found that the SD substitution enhanced the detection of the second binding event on interaction with BS69_CC-MYND_ (Figure 5C). The second binding event for T2 EBNA2_348-422_ SD was associated with a larger change in enthalpy (ΔH -7.65 kcal/mol) than the second binding event detected for T2 EBNA2_348-422_ (ΔH -1.59 kcal/mol) (Supplementary Table S2). Importantly we found that the affinities of the two binding events remained largely unaffected (Figure 5B and C), consistent with our earlier observations that the presence of serine at position 409 does not affect the ability of a type 2 polypeptide containing motif 1 and motif 2 to bind BS69 (Figure 1). To determine whether the impact of the SD mutation was dependent on the presence of BS69 binding motif 3, we also produced a polypeptide containing the SD and motif 3 mutation (EBNA2_348-422_ SD + m3 mt). We found that two binding events were still clearly detectable on interaction of this double mutant type 2 EBNA2 polypeptide with BS69_CC-MYND_ and that the enthalpy change of the second binding event was similar to that of the single SD mutant (ΔH -9.62 kcal/mol) (Supplementary Table S2) indicating that the impact of the SD change is still evident. The affinity of the second binding event (K_D2_) was however reduced by approximately 2-fold for EBNA2_348-422_ SD + m3 mt compared to the EBNA2_348-422_ SD mutant (Figure 5C and D). These data indicate that motif 3 contributes to the second binding event. We conclude that the SD mutation previously shown to enhance the growth maintenance properties of type 2 EBNA2 (21) does not affect BS69 binding but likely alters the conformation of the type 2 EBNA2 TAD. This may therefore impact on the binding of other transcriptional regulators that influence type 2 EBNA2 function.

### Full-length type 2 EBNA2 binds BS69_CC-MYND_ more efficiently in pull-down assays

To confirm our *in vitro* observations that a type 2 EBNA2 polypeptide binds an additional BS69 dimer, we examined the interaction of BS69_CC-MYND_ with full-length EBNA2 proteins stably expressed in B cells. Lysates from cells expressing type 1 or type 2 EBNA2 or the type 2 SD mutant were incubated with recombinant GST-BS69_CC-MYND_ immobilised on glutathione beads for increasing times and the amount of EBNA2 precipitated determined by Western Blotting (Figure 6). Consistent with the presence of an additional BS69 binding site in type 2 EBNA2, we found that GST-BS69_CC-MYND_ pulled down type 2 EBNA2 more efficiently than type 1 EBNA2 at short incubation times (Figure 6). In agreement with our *in vitro* observations using the type 2 EBNA2 SD mutant, we found that this protein interacted with BS69_CC-MYND_ with the same efficiency as type 2 EBNA2 (Figure 6). After 30 minutes incubation, GST-BS69_CC-MYND_ became saturated with EBNA2 and differences in association were no longer evident. A control GST fusion protein (GST-Rab11) did not precipitate EBNA2, confirming the specificity of the interactions. These data therefore confirm the increased association of BS69_CC-MYND_ with type 2 EBNA2.

**Figure 6.**
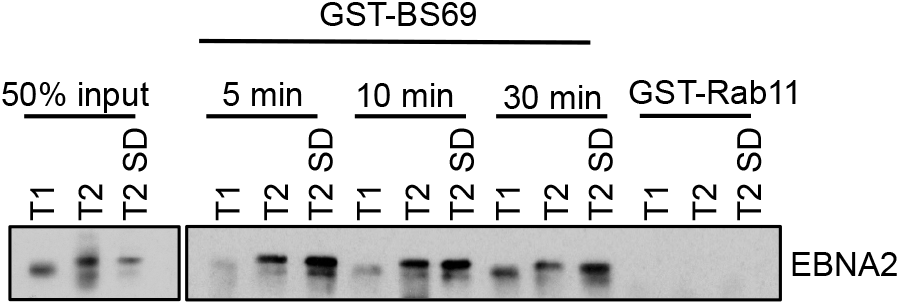
GST pulldown assay using GST-BS69_CC-MYND_ and lysates from B cell lines expressing type 1 and type 2 EBNA2. Nuclear extracts of Daudi:pHEBo-MT:EBNA-2 cells expressing type 1 (T1), type 2 (T2) or type 2 SD EBNA2 proteins were incubated for 5, 15 or 30 min at 4°C with glutathione beads which had been loaded with bacterial lysates expressing the GST-BS69_CC-MYND_. GST-RAB11B was used as a negative control and was incubated for 30 min with the nuclear extracts. Following washing, beads were resuspended in protein sample buffer and analysed by SDS-PAGE and Western blotting for EBNA2 using the PE2 anti-EBNA2 antibody. The EBNA2 proteins expressed by these cell lines display almost the same size (72 kD) since the number of polyproline residues had been equalised in all the EBNA2 alleles.

### Mutation of BS69 binding motif 3 in type 2 EBNA2 increases its growth maintenance function

To determine whether the presence of the additional BS69 binding motif in type 2 EBNA2 (motif 3) had functional consequences for the activity of type 2 EBNA2, we examined the ability of a type 2 EBNA2 motif 3 mutant to maintain B cell growth. We utilised a previously described assay using an EBV-infected LCL (EREB2.5) in which the activity of a type 1 estrogen receptor-EBNA2 fusion protein can be switched off by estrogen withdrawal (37). Loss of EBNA2 activity leads to growth arrest, but transfection of a stably-maintained plasmid expressing type 1 EBNA2 into these cells supports their survival (20). In contrast, the expression of type 2 EBNA2 cannot maintain the growth of these cells (20). We found that mutation of BS69 binding motif 3 produced a type 2 EBNA2 protein that was able to support the recovery of these cells from the loss of type 1 EBNA2 activity, with cells recovering well 2-4 weeks following estrogen withdrawal (Figure 7A). The type 2 EBNA2 motif 3 mutant behaved similarly to the type 2 EBNA2 SD mutant that was previously shown to support B cell growth in this assay (21). We also examined the ability of the SD and motif 3 double mutant in this assay and found that it showed a slightly increased ability to support B cells growth (Figure 7A). In our hands, expression of type 1 EBNA2 supported initial growth in this assay better than any type 2 mutants, with the mutants supporting growth recovery from 2 weeks (Figure 7A). We confirmed that all EBNA2 proteins were expressed at similar levels (Figure 7B). We conclude that the presence of the additional BS69 binding motif in type 2 EBNA2 impairs the ability of type 2 EBNA2 to maintain B cell growth.

**Figure 7.**
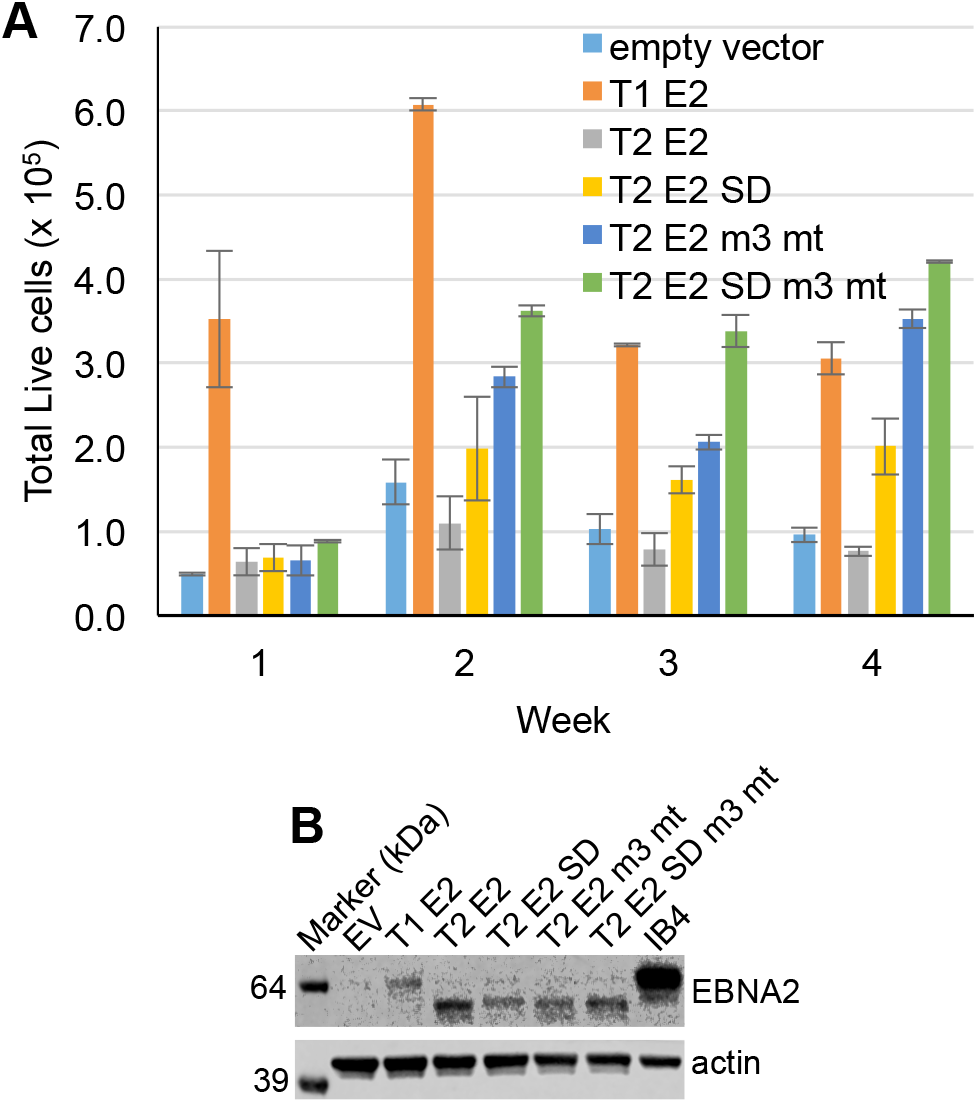
LCL growth maintenance assay using type 2 EBNA2 mutants. (A) ER-EB 2.5 cells conditionally expressing an estrogen receptor type 1 EBNA2 fusion protein were cultured in medium containing β-estradiol prior to resuspension in medium without β-estradiol. Cells were transfected with OriP-based plasmids (p294) expressing full length type 1 or type 2 EBNA2 or the type 2 SD, motif 3 or SD and motif 3 mutant EBNA2 proteins and cultured in medium without β-estradiol. Live cells (that excluded Trypan Blue) were counted 1, 2, 3 and 4 weeks post-transfection. Data from a representative experiment of 4 independent repeats is shown. Error bars show the mean ± standard deviation of duplicate cell counts for each sample. (B) Western blot analysis of EBNA2 expression in protein extracts from the transfected EREB2.5 cells. Cells were harvested 2 days after transfection. EBNA2 was detected using the PE2 monoclonal antibody and blots were probed for actin as a loading control.

### A BS69 isoform containing the MYND domain is expressed in type 1 and type 2 EBV-infected cells

BS69 functions as a negative regulator of EBNA2 transcription activity in reporter assays (34, 35), but previous studies have reported that BS69 expression is downregulated on infection of resting B cells by EBV and is low in the resulting immortalised LCLs (35). Transcriptional repression of BS69 by EBNA2 was implicated in BS69 downregulation indicating that EBNA2 may act to restrict expression of its own negative regulator (35). The cell lines examined in this previous study all harboured type 1 EBV or type 1 EBNA2, so we next addressed whether BS69 was expressed at similar levels in cells infected with type 1 and type 2 EBV. We examined BS69 protein levels in type 1 and type 2 LCLs using an anti-BS69 antibody raised against a region within the MYND domain of BS69. We found that BS69 was expressed at similar levels in type 1 and type 2 LCLs, but surprisingly levels in LCLs were similar to those in an EBV negative B cell line (AK31) (Figure 8A). We expanded our analysis to include additional EBV negative B cell lines (BJAB and DG75), EBV infected cell lines displaying the EBNA1 only latency I pattern of EBV gene expression (Akata and Mutu I), an additional type 1 LCL (IB4) and a BL cell line expressing all EBV latent proteins including EBNA2 (Mutu III) (both latency III cell lines) (Figure 8B). We found no correlation between BS69 expression and EBV infection or EBNA2 expression (Figure 8B). BS69 did not therefore appear to be downregulated as a result of EBNA2 expression. We also examined BS69 expression over the course of a primary B cell infection and found that BS69 was not downregulated as previously reported (Figure 8C).

**Figure 8.**
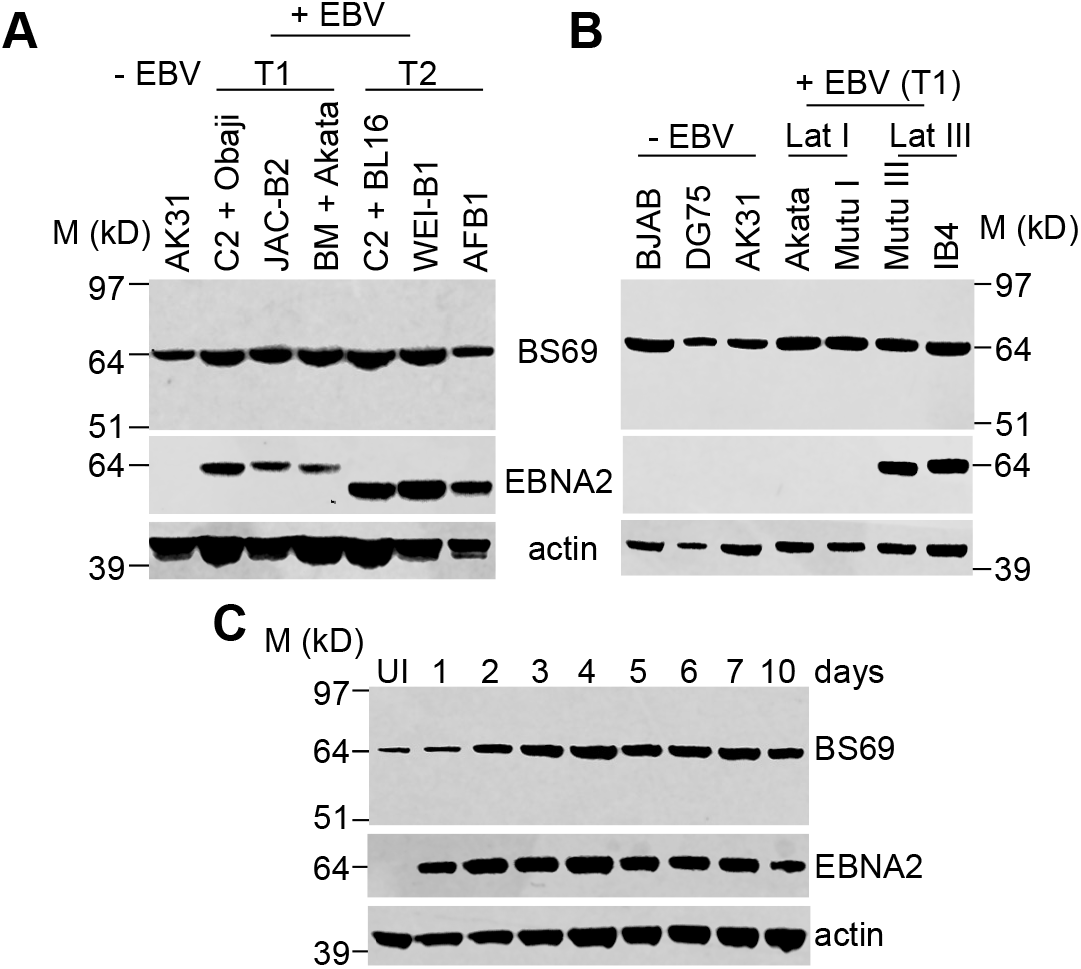
Western blot analysis of BS69 expression in EBV infected B cells. (A) Western blot analysis of BS69 expression in the EBV negative BL cell line AK31 and type 1 and type 2 EBV immortalised LCLs (that display the latency III pattern of gene expression associated with expression of all EBV latent proteins). BS69 was detected using an antibody that recognises a sequence in the MYND domain encoded by exon 15 (ab190890). EBNA2 was detected using the PE2 monoclonal antibody. Type 2 EBV has a lower molecular weight due a difference in the number of proline repeat residues present. Blots were probed for actin as a loading control. (B) Western blot analysis of BS69 expression in EBV negative (BJAB, DG75, AK31) and EBV positive latency I (Akata and Mutu I) and latency III B cell lines (Mutu III (BL) and IB4 (LCL) as in A. (C) Western blot analysis of BS69 expression on primary B cell infection. UI indicates uninfected primary B cells. Samples were taken at the indicated day post-infection and analysed as in A.

We therefore explored the possibility that we were detecting a different isoform of BS69. Alternative splicing has been reported to give rise to different BS69 isoforms and four have been experimentally verified (33) (Figure 9A). The canonical isoform (isoform 1, UniProt identifier: Q15326-1) contains 15 exons and encodes a protein of 71 kD (602 amino acids). Isoform 2 (Q15326-2) lacks amino acids 93-146 encoding the PHD domain (exon 4) and encodes a protein of 64.4 kD. Isoform 3 (Q15326-3) lacks amino acids 563-602 encoding the MYND domain (exon 15) and has a unique C-terminus encoded by an extended exon 14 sequence. Isoform 3 encodes a protein of 66.6 kD. Isoform 4 (Q15326-3) lacks exon 4 and exon 15 (and thus both the PHD and MYND domains) and encodes a protein of 60 kD. These isoforms were previously described as full length (FL), ΔPHD, ΔMD and ΔPHD, ΔMD respectively (33), but the exon numbering used in this previous study differed. The BS69 protein detected in Figure 8A and B has a molecular weight of approximately 64 kD consistent with that expected for isoform 2. This was the only protein detected by this antibody (against the MYND domain), indicating that isoform 1 was not expressed in the cell lines examined. Since the antibody we used would not detect BS69 isoforms 3 and 4, we could not exclude the possibility that one or more of these isoforms was also expressed and that an alternative BS69 isoform was detected previously (35). In line with this possibility, we noted that the QPCR analysis carried out by Harter *et al* used primers located in exon 4, which is absent from isoform 2.

**Figure 9.**
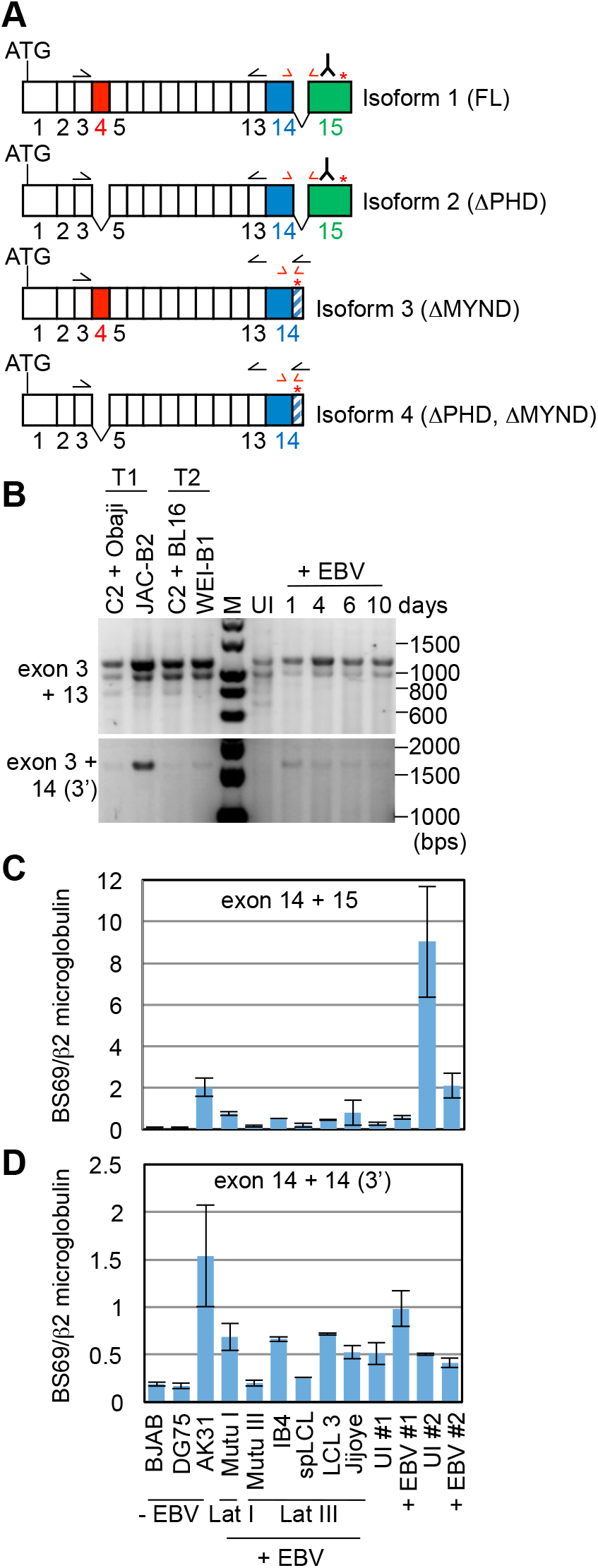
PCR analysis of the BS69 isoforms expressed in EBV infected B cells. (A) Diagram of the four experimentally verified BS69 isoforms. The position of the start codon (ATG) and stop codon (red asterisk) is indicated. Numbering of exons is *as per* the canonical isoform (isoform 1). Exon lengths are not to scale. Exon 4 is shown in red, Exon 15 is shown in green and exon 14 is shown in blue. The 3’ part of exon 14 present in isoforms 3 and 4 is shown in blue hatched lines. A sequence within exon 15 is recognised by the antibody (ab190890) used in Figure 8 as indicated on the diagram. Approximate locations of primers used for conventional PCR and QPCR are indicated by black and red arrows respectively. (B) Agarose gel analysis of PCR products generated using primers located in specific BS69 exons. The upper panel shows the PCR products amplified from cDNA samples from type 1 and type 2 LCLs and the primary infection samples shown in C using a forward primer located in exon 3 and a reverse primer located in exon 13. Transcripts containing exon 4 (isoform 1 and 3) will generate a 1139 bp product and those lacking exon 4 (isoforms 2 and 4) will produce a 977 bp product. The lower panel shows PCR carried out using the exon 3 forward primer and a reverse primer in the 3’ region of exon 14 that is uniquely present in differentially spliced BS69 transcripts lacking exon 15 (isoforms 3 and 4). Transcripts containing exon 4 and lacking exon 15 (isoform 3) will generate a 1578 bp product and those lacking exon 4 and exon 15 will produce a 1416 bp product (isoform 4). (C) QPCR analysis of cDNA from a panel of EBV negative and positive B cell lines and primary EBV infections using primers that amplify across the exon 14 and exon 15 junction. For EBV infections, samples from two different experiments (#1 and #2) were used. Primary B cells (uninfected, UI) were infected with EBV and samples harvested after 2 days (+ EBV). Results show the mean ± standard deviation of QPCR replicates from a representative experiment. BS69 relative quantities were normalized to β2 microglobulin. (D) QPCR analysis as in (C) using primers within exon 14. The reverse primer is located in the 3’ part of exon 14 only present in BS69 isoform 3 and 4.

No detail was provided on the anti-BS69 antibody used previously (35) and we were not able to find another antibody that detected isoform 3 and 4 in Western blotting. We therefore took a non-quantitative PCR approach to screen for different BS69 isoforms using cDNA prepared from LCLs and from B cells during a primary EBV infection. PCR using a forward primer in exon 3 and a reverse primer in exon 13 amplified two products indicating the presence of at least two different isoforms, one containing exon 4 (1139 bps) and one lacking exon 4 (977 bps) (Figure 9B). This would be consistent with the presence of isoform 3 (which contains exon 4) and isoform 2 (which lacks exon 4 and was detected by Western blotting (Figure 8)). PCR products were sequenced to confirm their identity (data not shown). However, since isoform 4 also lacks exon 4, this PCR analysis cannot rule out the additional presence of isoform 4. Since in isoforms 3 and 4 exon 15 is replaced by a short unique 3’ sequence from exon 14, we designed reverse PCR primers in this unique 3’ region. PCR using these primers amplified only one product of 1578 bps consistent with presence of exon 4 and the unique 3’ region (isoform 3) (Figure 9B). The identity of this PCR product was again confirmed by sequencing (data not shown). Importantly, we did not detect a smaller product (1416 bps) that would indicate the presence of isoform 4 (lacks exon 4 and exon 15). Our data therefore indicate that LCLs infected with either type 1 or type 2 EBV express both isoform 2 and isoform 3 of BS69.

To quantitatively examine whether either BS69 isoform 2 or isoform 3 were downregulated on EBV infection and in cells expressing EBNA2 as previously described (35), we used QPCR to analyse BS69 mRNA expression in primary B cells infected by EBV and in a panel of EBV negative and positive B cell lines. QPCR using primers that spanned exon 14 and exon 15 (present in isoform 1 and 2) detected variable levels of BS69 across the cell lines examined, with no obvious correlation with EBV positivity or EBNA2 expression (present in latency III EBV infected cell lines). This is consistent with the variability in BS69 protein expression detected in Western blot analysis of isoform 2 expression (Figure 8). Although, in one experiment (#2) primary B cells expressed high levels of BS69 isoform 2 mRNA that were reduced on EBV infection, the second primary infection experiment did not reproduce this observation. In fact, primary infection #2 was the same infection analysed by Western blotting in Figure 8C so this change in BS69 RNA expression did not result in decreased expression of BS69 isoform 2 protein. It is most likely therefore that BS69 isoform 2 expression varies in an EBV and EBNA2 independent manner. Analysis of BS69 mRNA expression using QPCR primers that would specifically amplify BS69 isoforms containing the long form of exon 14 (isoforms 3 and 4) also detected variable expression of BS69 that did not correlate with EBV positivity or EBNA2 expression indicating that isoform 3 expression is also EBV independent (Figure 9D).

We conclude that B cells infected with type 1 or type 2 EBV do not consistently display reduced expression of any detectable isoform of BS69 compared to uninfected B cells. Since BS69 isoform 2 contains the MYND domain that binds EBNA2 (that is absent in isoform 3), the continued expression of isoform 2 in EBV infected B cells would be expected to restrict the gene activation function of EBNA2.

### Inhibition of BS69 function increases EBNA2 transactivation activity

To determine whether inhibition of BS69 function increased EBNA2 transactivation function, we carried out EBNA2 transactivation assays in an EBV negative B cell line (BJAB) in which we overexpressed isoform 3 of BS69 lacking the MYND domain (ΔMYND) (but containing the coiled-coil dimerisation domain). This form of BS69 has been proposed to act as a dominant negative inhibitor of the MYND-domain dependent functions of BS69 (33). We performed transactivation assays using EBNA2-GAL4-DNA binding domain (DBD) fusion proteins and a Firefly luciferase reporter plasmid containing a synthetic promoter with 4 GAL4 binding sites. Plasmids expressing GAL4-DBD fusion proteins containing regions of type 1 EBNA2 (334-487) and type 2 EBNA2 (301-454) encompassing all BS69 binding motifs were transfected into BJAB cells in the presence or absence of plasmids expressing either full length BS69 (isoform 1) or isoform 3 (ΔMYND). Consistent with previous reports, we found that overexpression of full length BS69 inhibited transactivation by type 1 EBNA2 (34, 35) (Figure 10). BS69 also inhibited transactivation by type 2 EBNA2 (Figure 10). Consistent with its function as a dominant negative inhibitor, we found that expression of BS69 ΔMYND increased transactivation by both type 1 and type 2 EBNA2 (Figure 10). To determine whether this was a non-specific or EBNA2-dependent effect, we expressed BS69 ΔMYND in the absence of any GAL4-DBD-EBNA2 expressing constructs. In the absence of EBNA2 fusion protein expression, BS69 ΔMYND had no effect on the activity of the GAL4 reporter (Figure 10). These data therefore demonstrate that inhibition of the MYND-domain dependent function of BS69 in B cells relieves repression of EBNA2 transactivation. These data support our hypothesis that the expression of MYND-domain containing BS69 isoforms in B cells impedes EBNA2 gene activation function.

**Figure 10.**
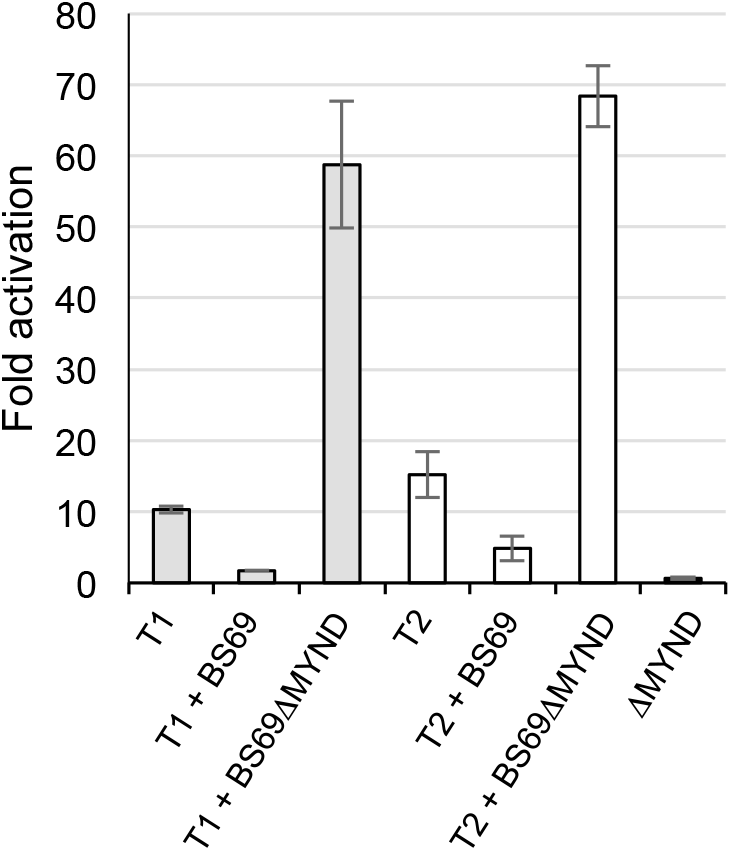
Expression of a dominant negative form of BS69 increases EBNA2 transactivation. Transactivation assays in BJAB cells using EBNA2-GAL4-DNA binding domain fusion proteins and a GAL4 reporter plasmid. Cells were cotransfected with 300 ng of either type 1 GAL4-DBD:EBNA2 (aa 334-487) or type 2 GAL4-DBD:EBNA2 (301-454) constructs, 500 ng of pFRLuc (Gal4 firefly luciferase reporter), 10 ng of pRL-CMV and 1 μg of BS69 (pCI-BS69) or BS69 ΔMYND (pCI-BS69-ΔMYND) expressing plasmids. For each sample, firefly luciferase values were normalised for transfection efficiency using Renilla luciferase values. Results are presented as luciferase activity relative to the pFR-Luc reporter plasmid plus empty vector (pcDNA3.1-GAL4-DBD). BS69 ΔMYND was also transfected in the absence of EBNA2 expressing constructs. Results show the mean of two independent experiments ± standard deviation.

Taken together our *in vitro* and cell-based assays suggest that during initial B cell infection the increased association of BS69 with type 2 EBNA2 may impede key gene activation events that are required for the efficient outgrowth of immortalised cell lines.

## Discussion

Type 2 EBV strains have reduced B cell transformation capacity and type 2 EBNA2 activates some viral and cell genes less efficiently than type 1 EBNA2, a feature that may underlie the impaired transformation phenotype. We have identified an additional binding site for the transcriptional repressor BS69 in the EBNA2 protein encoded by type 2 strains of EBV and show that mutation of this additional binding site improves the B cell growth maintenance properties of type 2 EBNA2. Our data therefore implicate increased BS69 association in the impaired function of type 2 EBNA2.

Type 2 EBV transforms resting B cells more slowly and results in the outgrowth of less immortalised cell clones than type 1 EBV (1, 19). Although early type 2 EBV transformants show reduced cell growth, the immortalised LCLs that eventually arise from a type 2 EBV infection grow similarly to those infected with type 1 EBV. Type 2 LCLs also maintain similar levels of expression of key EBNA2 target genes (20). This indicates that the impaired function of type 2 EBNA2 restricts an early stage in the B cell transformation process *in vitro*. Indeed two EBNA2 target genes that are only weakly activated by type 2 EBNA2 compared to type 1 EBNA2 (22), the viral oncogene LMP1 and the cell gene CXCR7, display slower and weaker induction during primary infection with type 2 EBV (20). Although it was previously reported that BS69 is downregulated on EBV infection, we found that there is continued expression of BS69 isoform 2 in EBV-infected cells. Since this isoform contains the MYND domain that mediates the BS69-EBNA2 interaction the expression of BS69 would be expected to restrict EBNA2 activation function. Consistent with this prediction we found that the expression of a dominant negative form of BS69 (isoform 3) lacking the MYND domain enhances EBNA2 activation function in B cells. Our data are consistent with a model where BS69 acts as a restriction factor for both type 1 and type EBNA2 but the association of type 2 EBNA2 with more molecules of the BS69 repressor protein further restricts the activation of growth and survival genes important in early transformation. Why a small number of specific genes are activated less well by type 2 EBNA2 is as yet not fully clear, but sequences resembling EICEs (bound by ETS and IRF transcription factors) are found at EBNA2 binding sites in the LMP1 promoter and binding sites closest to the cell genes that show reduced activation by type 2 EBNA2. Interestingly BS69 binding to PXLXP motifs in ETS2 has been shown to inhibit its transactivation activity (32). Since the ETS family member PU.1 is known to bind to the putative EICE in the LMP1 promoter and plays a role in LMP1 promoter activation, it is possible that type 2 EBNA2 functions less well in the context of PU.1 binding sites. Interestingly, PU.1 also contains PXLXP motifs that would be predicted to bind BS69, so enhanced tertiary complex formation between type 2 EBNA2, BS69 and PU.1 at the regulatory elements of specific genes may function to stabilise BS69 binding and further restrict gene activation by type 2 EBNA2.

Our data also provide important new molecular information on the nature of the complexes formed between EBNA2 and BS69 that may applicable to the way BS69 interacts with other cellular and viral transcription factors via its MYND domain. Single and multiple PXLXP BS69 binding motifs have been identified in the cell and viral binding partners of BS69, but the elucidation of the structure of the dimeric coiled-coil-MYND domain of BS69 led to a model that proposed that a BS69 dimer bound to the two adjacent PXLXP motifs (motif 1 and motif 2) in the same type 1 EBNA2 molecule (35). However, although the authors found that BS69 bound with increased affinity when two PXLXP motifs were present in EBNA2 polypeptides, the three-dimensional structure obtained comprised a BS69_CC-MYND_ dimer bound to two separate type 1 EBNA2 motif 1 peptides. Although the binding of motif 1 and motif 2 could be accommodated in the BS69_CC-MYND_ structure if the intervening 52 amino acids were looped out, the formation of this complex has not been formally demonstrated (35). Our SEC-MALS and SAXS analysis provides the first evidence that the BS69_CC-MYND_ dimer preferentially forms an intermolecular bridge between PXLXP motifs located on different EBNA2 molecules. This mode of binding is consistent with the fact that EBNA2 is a dimeric protein, with dimerisation mediated by the N-terminal END domain comprising amino acids 1-58 (36). Additional self-associating regions have also been mapped elsewhere in EBNA2 and include amino acids 97–121 and 122-344 (38, 39), although no molecular information is available on how these regions may contribute to dimerisation. Our data indicate that BS69 binding to sites in the C-terminal transactivation domain may contribute to the formation or stabilisation of EBNA2 dimers. Interestingly, although SEC analysis clearly demonstrated complex formation between both type 1 and type 2 EBNA 2 polypeptides and BS69_CC-MYND,_ the elution profiles of both complexes were broad. The type 1 EBNA2-BS69_CC-MYND_ elution profile had a clear shoulder indicating the presence of smaller MW complexes (Figure 3A). This would explain why the average MW determined by SEC-MALs was smaller than expected for a complex that contained two molecules of type 1 EBNA2 and two BS69_CC-MYND_ dimers. It is possible that in solution *in vitro* there is a mixed population of dimeric type 1 EBNA2 and monomeric type 1 EBNA2 complexes (where a single EBNA2 polypeptide is bound by one BS69_CC-MYND_ dimer as previously proposed). We were not able to investigate this further using SAXS as this ‘shoulder’ was not clearly defined, so SAXS analysis for both type 1 and type 2 EBNA2-BS69 complexes focused on the major elution peak of the large complex. Given that full length EBNA2 expressed in EBV-infected cells is a dimer, complexes involving two EBNA2 molecules are more likely to be physiologically relevant.

Surprisingly, in our GAL4-EBNA2 fusion protein assays we did not see weaker transactivation by the type 2 EBNA2 fusion protein compared to the type 1 EBNA2 fusion protein as reported previously (21). We used a longer region of EBNA2 compared to this previous study that encompassed all three BS69 binding sites for type 2 EBNA2 and the corresponding region of type 2 EBNA2 (with only two functional BS69 binding sites). Previously GAL4-EBNA2 fusion protein constructs were used that expressed a type 1 EBNA2 protein containing only BS69 binding motif 2 or the corresponding region of type 2 EBNA2 that contained BS69 binding motif 2 and 3 (21). It is not completely clear why the increased association of BS69 with type 2 EBNA2 is not associated with weaker transactivation in our assays in the context of a longer region of EBNA2, but it could point to the importance of the dimerisation that occurs in the context of the full-length protein in the assembly of larger BS69-EBNA2 complexes.

When considering the nature of assembly of BS69-EBNA2 complexes, it is likely that binding to motif 1 (which in type 1 EBNA2 has the highest affinity for BS69_CC-MYND_) would drive the initial interaction between EBNA2 and BS69 and binding to motif 1 probably constitutes the first binding event that can be distinguished in our ITC analysis using an increased number of injections. For type 2 EBNA2, since both motif 2 and 3 bind BS69 with similar affinity, binding to both of these motifs probably occurs with similar kinetics and is detectable as a single second binding event by ITC. Given the fact that BS69_CC-MYND_ dimers are predicted in the solution structure of the BS69-EBNA2 complex to be located side by side along a dimeric EBNA2 molecule, it is possible that interactions between BS69 coiled-coil dimers play a role in stabilising the oligomeric complex.

Our initial interest in examining type-specific binding of EBNA2 to BS69 centred around the influence of a serine residue in the TAD of type 2 EBNA2 that plays a key role in restricting B cell growth maintenance by type 2 EBNA2 (21). Although this residue is located immediately adjacent to BS69 binding motif 2 in type 2 EBNA2, we found that it did not increase BS69 binding (as might have been expected) when binding was compared to the corresponding region of type 1 EBNA2 where there is an aspartate residue in its place. It does not appear therefore that the influence of serine 409 on growth maintenance is mediated through alterations in BS69 binding affinity. Our ITC analysis however did find that a serine to aspartate change at this position in type 2 EBNA2 altered the nature of BS69 binding indicating that it may induce a conformational change in this region of EBNA2. This could result in differences in the binding of other transcription regulators to the type 2 EBNA2 TAD compared to the type 1 EBNA2 TAD. Possibilities could include increased binding of a repressor or co-repressor to the type 2 EBNA2 TAD or decreased binding of an activator or co-activator.

BS69 may have a wider role in regulating B cell transformation and the growth of EBV-infected cells in addition to its modulation of EBNA2 transactivation. BS69 localised to the cell membrane has also been implicated as an adaptor in signalling mediated by the EBV oncogene LMP1. The MYND domain of BS69 was reported to bridge an interaction between the carboxy terminal cytoplasmic domain of LMP1 and the TRAF6 signalling protein to activate the JNK signalling pathway (40). Conversely, BS69 has also been implicated as a negative regulator of LMP1-mediated NF-κB signalling by decreasing the association between C-terminal activation region (CTAR) 2 of LMP1 and the signalling adaptor TRADD (41) and by binding to CTAR1 and bringing in the negative regulator of NF-κB signalling, TRAF3 (42). Although further work appears to be required to fully understand the role of BS69 in LMP1 signalling and the relative proportions of nuclear and membrane-associated BS69, it is possible that BS69 is a key modulator of growth promoting events in EBV-infected cells. In this context, our work now sheds new light on how transformation by type 2 strains of EBV may be specifically curbed as a result of sequence variation that results in the creation of an additional binding site for BS69.

## Materials and Methods

### Cell lines

All cell lines were passaged twice weekly in RPMI 1640 media (Invitrogen) supplemented with 10% Fetal Bovine serum (Gibco), 1 U/ml penicillin G, 1 μg/ml streptomycin sulphate and 292 μg/ml L-glutamine at 37°C in 5% CO_2._ DG75 (43) and AK31 (44) are EBV negative BL cell lines and BJAB (45) is an EBV negative B cell lymphoma line. Akata (46) and Mutu I are EBV positive latency I BL cell lines and Mutu III is a cell line derived from Mutu I cells that drifted in culture to express all EBV latent proteins (latency III) (47). All LCLs also display the latency III pattern of EBV gene expression and were described previously (48); IB4, spLCL, LCL3, C2 + Obaji, JAC-B2, BM + Akata LCLs are infected with type 1 EBV and C2 + BL16, WEI-B1, Jijoye and AFB1 LCLs are infected with type 2 EBV. The ER-EB 2.5 LCL, expressing a conditionally active oestrogen receptor (ER)-EBNA2 fusion protein, was provided by Prof B. Kempkes and was cultured in the presence of β-estradiol (37). The Daudi:pHEBo-MT:E2T1, Daudi:pHEBo-MT:E2T2 and Daudi:pHEBo-MT:E2T2 S442D cell lines were generated and cultured as described previously (21). B cell infection samples were described previously (49).

### Plasmids

Constructs expressing N-terminal 6 x histidine tagged EBNA2 polypeptides were generated using the Sequence and Ligation Independent Cloning (SLIC) technique using type 1 EBNA2 (B95-8), type 2 EBNA2 (AG876) and type 2 EBNA2 SD (serine to aspartate at position 409) pSG5 expression plasmids as templates to amplify the regions of interest. DNA was PCR amplified using primers containing 20-30 bp of additional sequence from the regions 5’ and 3’ to the multiple cloning site of pET47b+. pET47b+ was digested using SmaI and HindIII and the PCR product and double-digested vector were then partially digested using the 3’ to 5’ exonuclease activity of T4 DNA Polymerase in the absence of dNTPs to generate long complementary 5’ overhangs. The PCR products and pET47b+ were then annealed on ice. The ligated DNA fragments obtained contain four nicks that are repaired by *E. coli* after transformation. For type 1 EBNA2, constructs encoded amino acids 381-445 or 381-455 to generate pET47b+ T1 EBNA2_381-445_ and pET47b+ T1 EBNA2_381-455._ Type 2 EBNA2 constructs encoded amino acids 348-412 or 348-422 to generate pET47b+ T2 EBNA2_348-412,_ pET47b+ T2 EBNA2_348-422_ and pET47b+ T2 EBNA2_348-422_ SD.

The BS69 CC-MYND domain (amino acids 480-602) was amplified from pCI-BS69 containing the full length human BS69 sequence (gift from Dr Stéphane Ansieau) and cloned using SLIC into the SmaI and HindIII sites of pET49b+ to generate a construct expressing an N-terminal GST-6x Histidine tag BS69_CC-MYND_ fusion protein.

To create GAL4-DNA-binding domain-EBNA2 TAD fusion protein expressing constructs pBlueScript plasmids carrying EBNA2 sequences were used as the template to PCR amplify the type 1 EBNA2 TAD (amino acids 426-463) and the type 2 EBNA2 TAD (amino acids 334-487) using *Taq* DNA polymerase. Primers contained *Bam*HI or *Not*I restriction sites at their 5’ ends. PCR products were first cloned into pCR2.1 using the TA cloning kit (Invitrogen) according to the manufacturer’s instructions. The pCR2.1 vector carrying the cloned PCR product was then digested with *Bam*HI and *Not*I and the EBNA2 TAD fragment then cloned into the *Bam*HI and *Not*I sites of pcDNA3.1-GAL4-DBD.

### Site-directed mutagenesis

Reverse PCR with the Q5^®^ Site-Directed Mutagenesis kit (NEB) was used to introduce mutations into BS69 binding motif 3 in the pET47b+ T2 EBNA2_348-422_ construct. This resulted in a change from PTLEP to ATAEA and generated pET47b+ T2 EBNA2_348-422_ motif 3 mt. The same primers were used to mutate motif 3 in the context of the SD mutation to generate pET47b+ T2 EBNA2_348-422_ SD + motif 3 mt.

### Growth maintenance assay

The EREB2.5 growth assay was performed as described previously (20). Briefly, 5 μg of OriP-p294 plasmids expressing type 1 EBNA2, type 2 EBNA2 or type 2 EBNA2 mutants were transfected into 5 × 10^6^ EREB 2.5 cells resuspended in 110 μl of buffer T using Neon transfection with 1 pulse of 1300 V for 30 msec. Following transfection, cell suspensions were added to 2 ml of media supplemented with 10% FBS and antibiotics but without β-oestradiol and incubated overnight in 12-well plates. The following day each transfected sample was made up to 10 mls with media and divided into 5 x 2 ml aliquots in a 12-well plate. Samples were harvested for cell counting and protein analysis at time points up to 4 weeks.

### Reporter assays

Cells were diluted 1:3 in fresh culture medium one day before transfection. 2 x 10^6^ BJAB cells were used for each individual transfection. Cells were pelleted by centrifugation at 335*g* for 5 minutes at 4°C and washed twice with pre-warmed PBS. Cells were resuspended in 100 μl of Neon resuspension solution R (Invitrogen). Cell suspensions were then mixed with plasmid DNA (2-12 μg in TE buffer). Cells were co-transfected with 300 ng of either type 1 GAL4-DBD:EBNA2 (aa 334-487) or type 2 GAL4-DBD:EBNA2 (301-454) constructs, 500 ng of pFRLuc (Agilent technologies), 10 ng of pRL-CMV (Promega) and 1 μg of BS69 (pCI-BS69) or BS69 ΔMYND (pCI-BS69-ΔMYND) expressing plasmids (gift from Dr Stéphane Ansieau). The DNA and cell mixture was transferred to a 100 μl Neon transfection pipette tip (Invitrogen). Cells were electroporated using Neon transfection protocol 14 (1200 V of pulse voltage, 20 ms of pulse width and 2 pulse number) and then transferred into 2 ml of pre-warmed media in a 6-well plate and incubated at 37°C for 24 hr.

Cell pellets were then lysed using 100 μl of 1X Passive Lysis buffer (Promega). Two freeze-thaw cycles were performed to achieve efficient lysis (20 sec on dry ice and thawing at room temperature for 2 min). Cell debris was removed by centrifugation at 25,000*g*, for 1 min at 4°C and the clear supernatant was then transferred to a fresh tube. 20 μl of lysate was assayed for firefly and Renilla luciferase activity using 20 μl of each dual luciferase assay kit reagent (Promega) and a microplate luminometer (LUMIstar Omega, BMG Labtech).

### Recombinant protein production

pGEX4T1-BS69 (kindly provided by Dr Stéphane Ansieau) was used to express a GST-BS69 fusion protein containing amino acids 452-602 of BS69 encompassing the CC-MYND domain (numbered according to the canonical isoform) (34). pGEX4T1-RAB11B expressing GST-tagged RAB11B (gift from Prof Gill Elliott) was used to produce a negative control protein for the GST pull-down assays.

For production of BS69_CC-MYND_ and EBNA2 polypeptides, the relevant plasmids were transformed into the Rosetta 2 (DE3) pLysS E. Coli strain and protein expression induced by adding 0.4 mM of isopropyl βD-1-thiogalactopyranoside (IPTG) to 3 litre cultures at an OD_600nm_ of 0.6. The bacteria were then grown at 20°C overnight before harvesting for protein purification. Cell pellets from a 3 litre induced culture were lysed for 30 minutes on ice with constant stirring in 100 ml of lysis buffer (25 mM Tris-HCL pH 7.5, 500 mM NaCl, 5% Glycerol). The lysis buffer was supplemented with 0.25 mg/ml lysozyme, 2 mM MgCl_2_, 1 mM TCEP (tris(2-carboxyethyl)phosphine), two protease-inhibitor complete tablets (Roche) and DNase and 0.2 mg/ml of DNase I. Lysates were then sonicated at 37% amplitude for 5 minutes with 10 seconds pulses using a Vibra-cell sonicator (SONICS). The cell debris was pelleted at 15000 rpm for 45 minutes at 4°C (Biofuge Stratos, Heraeus). Beads from 3 ml of HisPur™ Cobalt Resin slurry (Thermo Scientific) were added to the cleared lysate along with 2 mM imidazole and the sample incubated for 1 hour and 30 minutes at 4°C with rolling. Samples were decanted into a centrifuge column (Thermo Scientific Pierce) and washed with buffer (25mM Tris-HCl, 500mM NaCl and 1mM TCEP, 2 mM of imidazole, pH 7.5). The protein was eluted from the beads using buffer containing increasing concentrations of imidazole (5 mM, 10 mM, 20 mM, 50 mM, 100 mM, 200 mM, 300 mM and 500 mM). Fraction samples were analysed by SDS-PAGE and the fractions containing recombinant protein were pooled and incubated with 3C protease (200 μl of 2 mg/ml) in the presence of 2 mM DTT overnight at 4°C to cleave the Histidine tag. His-tag cleaved proteins were then separated from the 3C protease by passing the protein sample through a GSTrap HP column (Amersham) using a peristaltic pump to capture the GST-tagged 3C protease. Untagged recombinant proteins were then concentrated and injected into a HiLoad™ 16/600 Superdex™ 75g Column (GE Healthcare) pre-equilibrated in 25 mM Tris-HCL, 200 mM NaCl, 1 mM TCEP, pH 7.5 purified at 0.5 ml/min. Protein fractions containing purified protein were then pooled, concentrated and stored at -80°C until required. Approximately 1 mg of purified EBNA2 polypeptides or 4 mg of BS69_CC-MYND_ was obtained from 1 litre of culture.

### Isothermal titration calorimetry

Four commercially synthesized peptides (Peptide Synthetics) were used for ITC. These included BS69 binding motif 2 of type 1 EBNA2 (435-445) or type 2 EBNA2 (402-412) and putative BS69 binding motif 3 of type 1 EBNA2 (445-455) or type 2 EBNA2 (412-422). Frozen protein was quickly thawed using running water and dialysed overnight at 4°C using Slide-A-Lyzer^®^ MINI Dialysis Units (Thermo Scientific) against ITC buffer (20mM Tris-HCl, 100mM NaCl and 1mM TCEP, pH 7.5). The next day, protein samples were centrifuged at 13000 rpm for 10 minutes at 4°C and the concentration was determined by NanoDrop spectroscopy (NanoDrop Technologies) with their respective molecular weights and extinction coefficients. EBNA2 peptides (1mM) and polypeptides (type 1, 0.3 mM and type 2, 0.6 mM) were titrated against BS69_CC-MYND_ (0.1mM) at 25°C using a MicroCalTM iTC200 instrument (Malvern). For peptides, 13 x 3.0 μl injections were used for titration. For EBNA2 polypeptides 19 x 2.0 μl or 29 x 1.3 μl injections were used for titration. ITC data were corrected for non-specific heat and analysed using MicroCal Origin^®^ 7.0. The experiments were performed in triplicate alongside a control experiment with no BS69_CC-MYND_ (buffer only in the cell). All polypeptides were used within 24 hours of dialysis into ITC buffer.

### Size Exclusion Chromatography

An S200 10/300 GL gel filtration column (GE Healthcare) was equilibrated with buffer containing 20mM Tris-HCl, 100mM NaCl and 1mM TCEP, pH 7.5. Individual EBNA2 or BS69_CC-MYND_ polypeptides or complexes were applied to the column and analysed at a flow rate of 0.5ml/min. The eluted fractions were then analysed by SDS-PAGE and Quick Coomassie staining.

### Size Exclusion Chromatography-multi-angle light scattering

EBNA2-BS69 complexes were prepared by pre-incubating proteins in a 1:3 molar ratio for at least 30 mins at 4°C. Purified samples (45 μl) at a concentration of 5 mg/ml were loaded onto a Shodex KW403-4F column at 25°C pre-equilibrated in 20 mM Tris-HCl, 100mM NaCl and 1mM TCEP, pH 7.5. Elution fractions were monitored using a DAWN HELEOS II MALS detector followed by a refractive index detector Optilab T-rEX (Wyatt Technology). Molecular masses of each individual peak were determined using ASTRA 6 software (Wyatt Technology). For normalization of the light scattering and data quality, BSA was used as a calibration standard.

### Small-angle X-ray scattering

Synchrotron radiation X-ray scattering data from solutions of individual proteins or complexes prepared as for SEC-MALS were collected on beamline B21 at Diamond Light Source (Didcot, United Kingdom), with an inline HPLC system. X-ray scattering patterns were recorded on a Pilatus detector after injection of 45 μl of protein sample (5-10 mg/ml) in a Superdex 200 3.2/300 column equilibrated in 20mM Tris-HCl, 100mM NaCl, 2% Sucrose and 1mM TCEP, pH 7.5. Samples were analysed at 20°C using a flow-rate of 0.25 ml/min. Initial data processing (background subtraction and radius of gyration Rg calculation) was performed using ScÅtter (v3.0 by Robert P. Rambo; Diamond Light Source). *Ab initio* beads model for the complex were prepared using DAMMIF (50). 23 independent dummy atom models were obtained by running the program in ‘slow’ mode. DAMAVER was then used to align and average the models (51). The *ab initio* generated beads models were refined using DAMMIN and compared to the experimental scattering data to derive χ^2^ values (52). The goodness-of-fit χ^2^values for the docked structure compared to the experimental scattering data were determined with FoXS (53).

### GST pull-down assays

Nuclear extracts were prepared from control or EBNA2 expressing Daudi cell lines. EBNA2 expression in Daudi:pHEBo-MT:EBNA2 cells was induced with 5 μM CdCl_2_ for 24 hours. At least 4×10^7^ cells were then harvested and resuspended in 1 ml of buffer A (10 mM HEPES pH 7.9, 1.5 mM MgCl2, 10 mM KCl, 0.5 mM 1,4-dithiothreitol (DTT) (Sigma), 1 mM PMSF (Sigma) and 1x complete protease inhibitor cocktail (Roche)). Cells were pelleted by centrifugation at 1000*g* for 5 min at 4°C and lysed in 100 μl of buffer A supplemented with 0.1% (v/v) NP-40 and incubated on ice for 5 min. Cell lysates were centrifuged at 2700*g* for 30 sec at 4°C and the nuclei resuspended in 50 μl of buffer B (20 mM HEPES pH 7.9, 420 mM NaCl, 1.5 mM MgCl2, 0.2 mM EDTA, 1 mM PMSF, 25% (v/v) glycerol, 1 mM DTT and 1x complete protease inhibitor cocktail) at 4°C for 20 min with rotation. Samples were finally centrifuged at 11,600*g* for 10 min at 4°C and the supernatants/nuclear extracts were transferred to fresh eppendorf tubes and the protein concentration was determined before storage at -80°C.

Lysates containing GST-tagged BS69_CC-MYND_ or GST-RAB11B were prepared from 100 ml cultures of E. Coli BL21 (DE3). Transformed cells were cultured at 30°C until they reached an OD_600nm_ of 0.6 and protein expression was induced at 25°C with 0.5 mM IPTG for 3-4 h. Cells were pelleted at 2,800*g* for 20 min at 4°C and then resuspended in 10 ml of Lysis buffer (20 mM Tris-Cl pH 8.0, 150 mM NaCl and 1 mM DTT) supplemented with 120 μl of lysozyme (10 mg/ml) and lysates incubated on ice for 20 min. Lysates were sonicated at high speed for 3 x 15 sec pulses in ice water using an Ultrasonic XL2020 Processor (Heat Systems) and cell debris pelleted at 17,900*g* for 30 min at 4°C. Lysates were stored at -80°C until required.

For pull-down assays, 50 μl of 50% Glutathione-Sepharose 4B Bead slurry (GE Healthcare) was washed three times in ice-cold binding buffer (20 mM Tris-Cl pH 8.0, 150 mM NaCl, 1 mM DTT and 0.1 mg/ml BSA). Beads were pelleted by centrifugation at 25,000*g* for 1 min and 100 μl of bacterial lysate containing the GST-tagged protein was incubated with the washed beads for 1 h at 4°C with rotation. Glutathione-Sepharose Beads bound to the GST-tagged protein were then washed with ice-cold binding buffer six times and pelleted by centrifugation at 25,000*g* for 1 min. Loaded GST-BS69_CC-MYND_ beads were then incubated with nuclear extracts containing EBNA2 at 4°C for different times (5, 10 and 30 minutes). Loaded GST-RAB11B beads were incubated with lysates for 30 minutes. Beads were then washed six times with ice-cold binding buffer and pelleted by centrifugation at 25,000*g* for 1 min. Beads were then resuspended in 25 μl of 2x SDS sample buffer (120 mM Tris-Cl pH 6.8, 4% (w/v) SDS, 2% (v/v) β-mercaptoethanol, 20% (v/v) glycerol and 0.01% (w/v) bromophenol blue) and incubated at 95°C for 5 min and analysed for EBNA2 levels by SDS-PAGE and Western blotting.

### SDS-PAGE and Western blotting

SDS-PAGE and Western blotting was carried out as described previously (54, 55) using the anti-EBNA2 monoclonal antibody PE2 (gift from Prof M. Rowe) anti-actin 1/5000 (A-2066, Sigma) and anti-BS69 1/1000 (ab190890, Abcam). Western blot visualisation and signal quantification was carried out using a Li-COR Imager. Gels were stained using Quick Coomassie stain (Generon Ltd).

### PCR and QPCR

RNA was extracted from cells using Trireagent (Sigma), further purified using the RNeasy kit (Promega) and cDNA synthesised using random primers and the ImProm II reverse transcription kit (Promega). Standard PCR reactions were performed with Phusion DNA polymerase (New England Biolabs) using the relevant BS69 primers listed in Supplementary Table S1. Quantitative PCR was performed in duplicate using the standard curve absolute quantification method on an Applied Biosystems 7500 real-time PCR machine as described previously (56) using the relevant primers listed in Supplementary Table S1. The efficiency of all primers was determined prior to use and in each experiment and all had amplification efficiencies within the recommended range (90–105%).

## Data Availability

SAXS data have been deposited in the small angle scattering biological data bank (SASBDB) (www.sasbdb.org) under accession numbers SASDEF6, SASDEG6, SASDEH6, SASDEJ6, SASDEK6;

https://www.sasbdb.org/data/SASDEF6/5llm1wasc4/

https://www.sasbdb.org/data/SASDEG6/4yff1ro01u/

https://www.sasbdb.org/data/SASDEH6/tpia09j2m0/

https://www.sasbdb.org/data/SASDEJ6/werm0avk55/

https://www.sasbdb.org/data/SASDEK6/07t9uflv7k/

## Acknowledgements

We thank Dr Stéphane Ansieau and Prof Gill Elliott for providing plasmids. We thank Diamond Light Source for beamtime (proposal mx14891) and the staff of beamline B21 for assistance with data collection.

## Supporting Information Legends

**Supplementary Figure S1.** Normalised (dimensionless) Kratky plots generated using ScÅtter (v3.0 by Robert P. Rambo; Diamond Light Source). I(q)/I(0)*(q*Rg)2 vs q*Rg. Scattering intensity I (q), scattering vector (q), radius of gyration (Rg).

**Supplementary Figure S2.** Solution structures of BS69_CC-MYND_ and type 1 and type 2 EBNA2 polypeptides determined by SAXS. (A-C) The SAXS envelopes (grey mesh) were generated by averaging 20 *ab-initio* models using the DAMMIF programme and further refined with DAMMIN to produce refined dummy atom models (magenta mesh). The maximum dimension (D_max_) and volume were calculated using the ScÅtter programme. In (A) the BS69_CC-MYND_ dimer structure (cyan; PDB ID: 5HDA) was manually docked into the envelope. (D-F) SAXS scattering data (black dots) fitted to the *ab initio* DAMMIN dummy atom (red line). χ^2^ values for fitting are shown.

**Supplementary Figure S3.** Alternative models and their respective goodness-of-fit to the experimental SAXS data. (A) One BS69_CC-MYND_ dimer (cyan; PDB ID: 5HDA) and the *ab initio* model of one type 1 EBNA2_381-455_ polypeptide (salmon) were manually docked into the *ab initio* envelope (grey mesh) of the type 1 EBNA2 BS69 complex. (B) SAXS scattering data were fitted to the docked structural complex shown in A and gave a χ^2^ of 13.33 using FoXS. Graphs show relative log intensity vs scattering vector (q) (upper panel) and the deviation (residual) of the model from the experimental data (lower panel). The hydration parameter (C_2_) was fixed to 0 to prevent the hydration shell increasing to beyond the maximum limit of 4 to attempt to fit the structure into the envelope. (C) The structural model shown in Figure 4C (two BS69_CC-MYND_ dimers and two type 1 EBNA2_381-455_ polypeptides) was refitted to the SAXS envelope using FoXS with the C_2_ value set to 0 for comparison. This gave a similar χ^2^ (2.59) to that shown in Figure 4A indicating a much better fit to the scattering data. (D) Two BS69_CC-_ MYNDBS69_CC-MYND_ dimers (cyan; PDB ID: 5HDA) and the *ab initio* model of one type 2 EBNA2_348-422_ polypeptide (orange) were manually docked into the *ab initio* envelope (grey mesh). (E) SAXS scattering data were fitted to the docked structural complex shown in D and gave a χ^2^ of 4.65 using FoXS. Graphs show relative log intensity vs scattering vector (q) (upper panel) and the deviation (residual) of the model from the experimental data (lower panel). The hydration parameter (C_2_) was fixed to 0 to prevent the hydration shell increasing to high levels to attempt to fit the structure into the envelope. (E) The structural model shown in Figure 4D (three BS69_CC-MYND_ dimers and two type 2 EBNA2_348-422_ polypeptides) was refitted to the SAXS envelope using FoXS with the C_2_ value set to 0 for comparison. This gave a similar χ^2^ (1.47) to that shown in Figure 4B indicating a much better fit to the scattering data.

**Supplementary Table S1**. Primer sequences.

**Supplementary Table S2.** Data obtained from Isothermal calorimetry analysis of EBNA2 peptides and polypeptides binding to BS69_CC-MYND_ (n.d. indicates binding not detected). Data show the mean ± standard deviation for three independent experiments. For peptides n values were fixed to 1. Data from type 2 EBNA2 peptides and polypeptides are in shaded columns.

**Supplementary Table S3.** SAXS data for BS69 and EBNA2 polypeptides and complexes.

